# Fly-CURE, a Multi-institutional CURE using *Drosophila*, Increases Students’ Confidence, Sense of Belonging, and Persistence in Research

**DOI:** 10.1101/2023.01.16.524319

**Authors:** Julie A. Merkle, Olivier Devergne, Seth M. Kelly, Paula A. Croonquist, Cory J. Evans, Melanie A. Hwalek, Victoria L. Straub, Danielle R. Hamill, David P. Puthoff, Kenneth J. Saville, Jamie L. Siders, Zully J. Villanueva Gonzalez, Jackie K. Wittke-Thompson, Kayla L. Bieser, Joyce Stamm, Alysia D. Vrailas-Mortimer, Jacob D. Kagey

## Abstract

The Fly-CURE is a genetics-focused multi-institutional Course-Based Undergraduate Research Experience (CURE) that provides undergraduate students with hands-on research experiences within a course. Through the Fly-CURE, undergraduate students at diverse types of higher education institutions across the United States map and characterize novel mutants isolated from a genetic screen in *Drosophila melanogaster*. To evaluate the impact of the Fly-CURE experience on students, we developed and validated assessment tools to identify students’ perceived research self-efficacy, sense of belonging in science, and intent to pursue additional research opportunities. Our data show gains in these metrics after completion of the Fly-CURE across all student subgroups analyzed, including comparisons of gender, academic status, racial and ethnic groups, and parents’ educational background. Importantly, our data also show differential gains in the areas of self-efficacy and interest in seeking additional research opportunities between Fly-CURE students with and without prior research experience, illustrating the positive impact of research exposure (dosage) on student outcomes. Altogether, our data indicate that the Fly-CURE experience has a significant impact on students’ efficacy with research methods, sense of belonging to the scientific community, and interest in pursuing additional research experiences.

## INTRODUCTION

As undergraduate STEM education continues to evolve and make improvements that facilitate the training of scientists from diverse backgrounds, it is becoming increasingly apparent that an authentic research experience is key for promoting student persistence within STEM majors and for adequate preparation for future scientific careers. There has been a national call for all STEM majors to have such an experience during their undergraduate education (1, 2), however, a significant challenge to this call is simple logistics. While some undergraduates do participate in a traditional apprentice-based research experience, there is not enough research lab capacity to accommodate all undergraduate STEM majors (3). One response to limited research opportunities has been to incorporate authentic research experience(s) into the curriculum. Such courses, often referred to as CUREs (Course-based Undergraduate Research Experiences), provide a research experience to a larger number of students (approximately 20-25 students per faculty or teaching assistant mentor) within a single iteration (3–5). Several CURE-type endeavors have been developed and, consequently, have provided research opportunities to a far greater number of STEM undergraduates than would have been possible through mentored bench research alone (5– 11).

CURE participation positively impacts science education in several ways. In comparison with traditional apprenticeships, CUREs not only reach more students, but also represent a more inclusive approach to research (3, 12). Student participation in CUREs has been shown to enhance critical thinking skills (10, 13), increase learning gains, bolster scientific identity (14, 15), and increase interest in science and scientific research (16). Each of these outcomes is likely an important factor driving the positive correlation between student participation in CUREs and increased STEM retention rates, including for underrepresented minority students (11, 17, 18).

Faculty, departments, and the scientific community at large can also be positively impacted by implementing CURE pedagogies. Faculty at Primarily Undergraduate Institutions (PUIs) typically have a heavy teaching requirement (teaching 3-4 classes per semester is not uncommon) that often comes with the additional expectation of research productivity (19). CUREs provide such faculty with an opportunity to combine teaching and research into a single endeavor that can, when properly structured and implemented, produce publishable work (both research data collected/analyzed by the students and pedagogical data measuring the impact on students) (16, 17, 20, 21). However, setting up a successful CURE comes with many challenges, the largest of which is typically the identification of a research project that is feasible for undergraduates working within the confines of a laboratory course (meeting 1-2 times per week, 3-5 hours total), budget-friendly, and longitudinally sustainable. The implementation of CUREs by regional and national consortia has been successful in overcoming many of these challenges. Efforts such as Science Education Alliance (SEA-PHAGES), Genomics Education Partnership (GEP), and Small World Initiative, have had success with CURE implementation at multiple sites, due in part to offering established, ready-to-go projects that entice faculty participation by reducing the burden of identifying a suitable research project and developing the infrastructure to support these projects (22–24). Not only does this approach provide research opportunities for more students, but it also increases the amount of valuable undergraduate-generated data. In addition, faculty and student participants are typically authors on research papers that include their contributing data. Here we describe a new CURE consortium called Fly-CURE that utilizes *Drosophila melanogaster* as a research model in undergraduate genetics laboratory courses.

The Fly-CURE was established in 2012 at the University of Detroit Mercy and centers on characterizing and mapping novel EMS-induced mutations isolated in a genetic screen for genes that regulate cell growth and cell division within the developing *Drosophila* eye (25). In the Fly-CURE, students start with an uncharacterized mutant and, in its analysis, learn about and utilize a variety of techniques commonly taught in more traditional undergraduate genetics laboratory courses. The Fly-CURE curriculum includes, but is not limited to classical Mendelian genetics, molecular genetics, and bioinformatics. Over the last ten years, students participating in the Fly-CURE have characterized over twenty novel *Drosophila* mutations, which have been published in eleven publications and included 581 student co-authors (26–36). Currently, the Fly-CURE is being taught at over twenty institutions across the United States. The institutional diversity of the Fly-CURE consortium has allowed us to measure the impact of the Fly-CURE pedagogy on a variety of student attitudes, including their sense of belonging in STEM, research competency, and intent to continue toward a STEM career. We also evaluated the effect of dosage on these metrics, where dosage refers to research experiences that a student participated in prior to participation in the Fly-CURE research project. Assessing the impact of research experience “dosage” on STEM undergraduates participating in the Fly-CURE consortium may shed light upon whether there is a critical number of research experiences that impact students’ retention and ultimate success in STEM fields.

## METHODS

### Fly-CURE consortium: institutions, faculty, and student participants

Pre- and post-survey data were gathered from 480 Fly-CURE students over three academic years: 2019-2020, 2020-2021, and 2021-2022. The demographics of the participating schools and students are detailed in Appendix 1 and shown in Figure 2. In the years of data collection and in the data presented, there were 15 faculty who implemented the Fly-CURE across 14 institutions.

**Figure 1.**
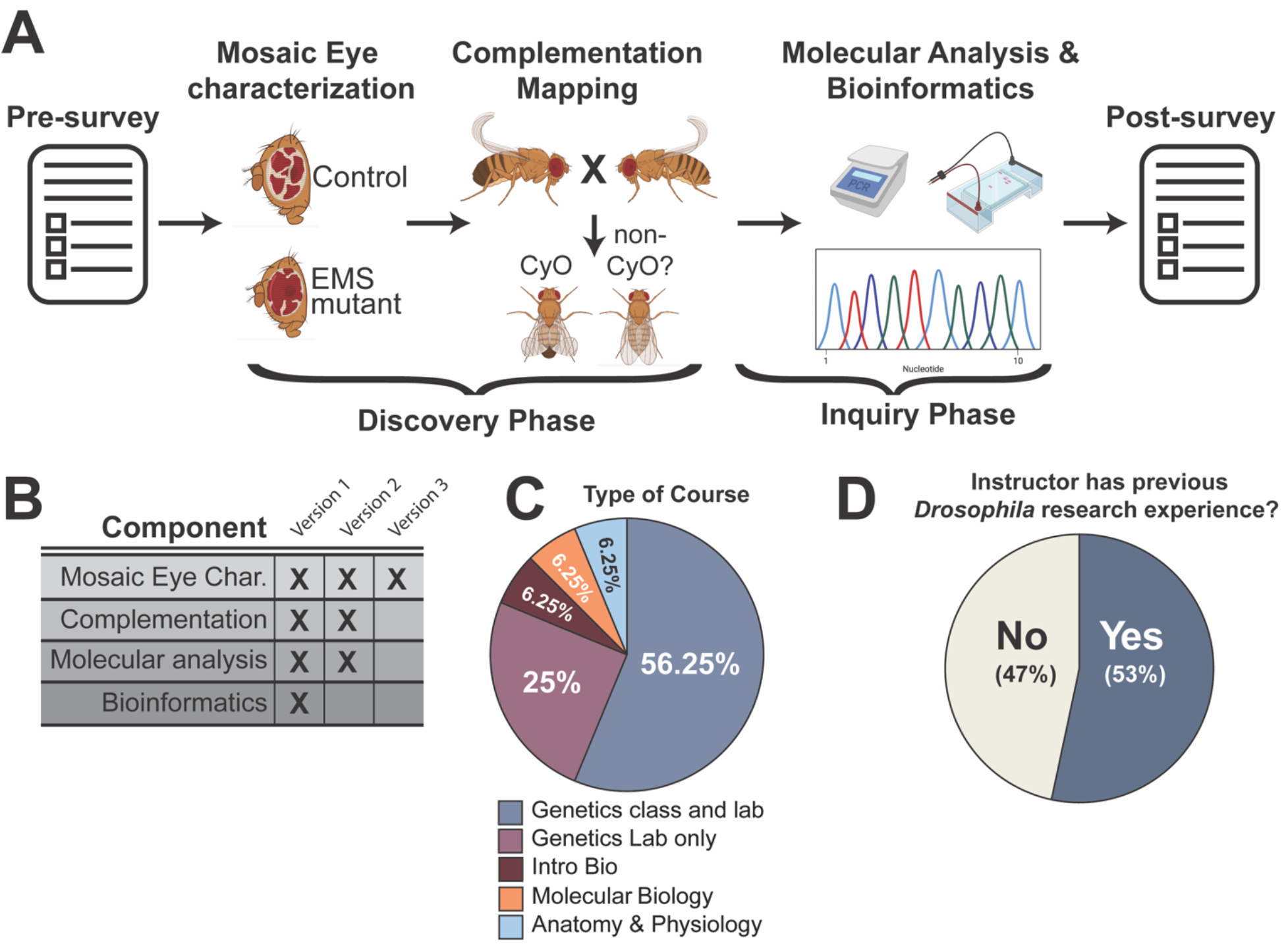
The Fly-CURE is a modular course-embedded research project. (A) Students enrolled in the Fly-CURE took an initial survey in which students reported their perceived self-efficacy in research and sense of belonging in science. The pre-course survey was also used to collect student demographic information. An FRT/Flp-based approach was used to create mitotic clones in *Drosophila* eye tissue where tissue homozygous for an EMS-induced mutation was marked by red pigment and wild-type tissue was marked by the absence of eye pigment. The growth ability of tissue homozygous for the EMS mutation was assessed by comparing the amount of red (mutant) to white (wild type) tissue within the adult fly eye. In parallel, the genomic locus of the mutation on chromosome 2R was then determined by complementation mapping with defined chromosome deletions. Once this initial “discovery” phase was completed, students initiated a more hypothesis-driven “inquiry” phase of the project. Bioinformatics and molecular approaches were used to design PCR primers and then amplify and sequence a portion of the chromosomal region that fails to complement the mutation. Finally, a post-course survey was implemented to measure the impact of the Fly-CURE on students’ perceived self-efficacy in research, sense of belonging in science, and intent to pursue additional research experiences or scientific careers. (B) Different combinations of the Fly-CURE components can be combined in a modular format, depending on the learning objectives of the course where the Fly-CURE was implemented (also see Appendix 1). (C) While most courses implementing the Fly-CURE were genetics courses with a lab or a stand-alone genetics lab course, the Fly-CURE was incorporated into a variety of other undergraduate Biology courses (Appendix 1). (D) 53% of Fly-CURE instructors (8 out of 15) had previously worked in a research setting using *Drosophila melanogaster*.

**Figure 2.**
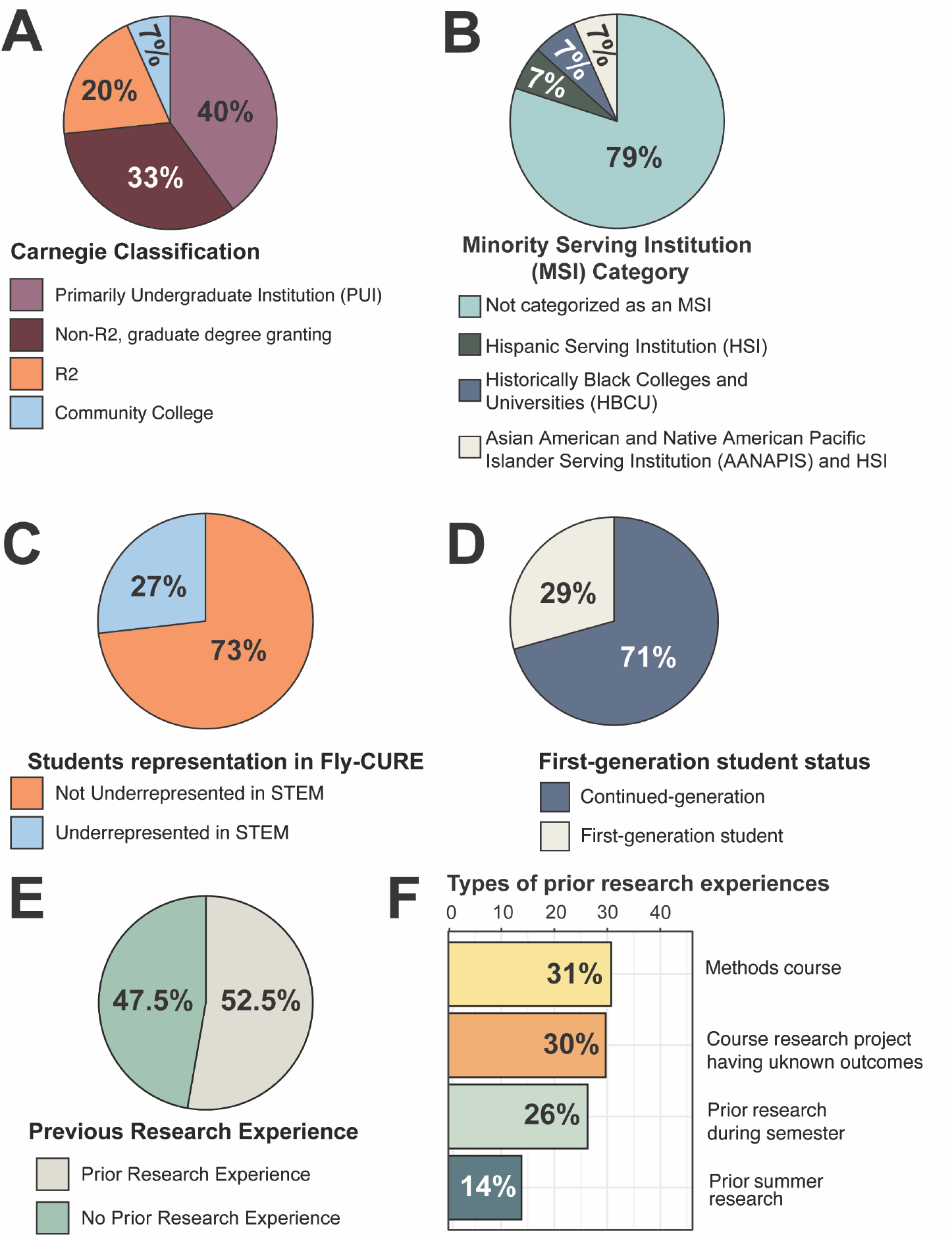
Institutional, demographic, and previous research experience of students enrolled in the Fly-CURE. (A) Institutional profiles where the Fly-CURE was implemented were obtained from The Carnegie Classification system. Institutions classified as Baccalaureate Colleges were combined into a single Primarily Undergraduate Institution (PUI) category. Carnegie Institutions classified as Doctoral/Professional Universities or Master’s Universities were pooled together as Non-R2, graduate degree-granting institutions. Number of institutions in each category: PUI (n=6), Non-R2 graduate degree-granting institutions (n=5), R2 (n=3), Community College (n=1) (see Appendix 1). (B) Minority Serving Institution (MSI) data was obtained from The Office of Postsecondary Education Eligibility Matrix. Number of institutions in each category: Non-MSI (n=12), Hispanic Serving Institution (HSI, n=1), Historically Black College or University (HBCU) (n=1), Asian American and Native Pacific Islander Serving Institution (AANAPIS) and HSI (n=1) (Appendix 1). (C-F) Demographic information from the student pre-course survey was used to determine the number of students that self-identified as underrepresented in STEM (C) or as first-generation college students (D). Pre-course survey data was also used to identify whether Fly-CURE participants had previously obtained research experience (E) and if so, the type of research experience in which students had participated (F).

All participating students were asked to complete a voluntary online survey before beginning (pre-course) and after completing (post-course) a Fly-CURE course offering (see Figure 1A for workflow). Approval to assess students was obtained by each participating institution from their Institutional Review Board. After each semester, responses were collected and analyzed by SPEC Associates (Southfield, MI), an independent analytics firm specializing in education and learning. Confidentiality was maintained by providing each instructor with a unique link to the online surveys that could be distributed to students. SurveyMonkey was the online platform used, with completed surveys being directly received by SPEC Associates without the instructors’ ability to see responses. The components of the pre- and post-course surveys used for this study are available in Appendix 2.

From the 895 students invited to participate in the surveys, we received 740 completed pre-course surveys and 683 completed post-course surveys. Only data in which students took both the pre- and post-course surveys were used in our analysis (69% of surveys were pre-/post-test matched). Pre- and post-survey responses were matched based on answers to non-identifying questions such as childhood home address. Student attentiveness was also assessed using one inattentive item on both the pre- and post-survey. Students who did not respond accordingly were eliminated from the analysis. Ternovski and Orr provide evidence that survey respondents who are inattentive also provide less reliable demographic data and are systematically different from attentive respondents (37). Following analysis for student attentiveness and pre-/post-test pairing, 65% of surveys were included in our current study. The number of surveys used in each comparative analysis differed because some students responded to only a subset of the survey items.

Participants identified their gender as female (69%), male (28%), their gender was not listed (1%), or they preferred not to say (2%). Participants were from ethnic or racial groups classified by the NSF as underrepresented in STEM (27%) and groups not considered underrepresented in STEM (73%). Demographic groups who were considered underrepresented in STEM were the following: Native Hawaiian or other Pacific Islander (original peoples), American Indian or Alaskan Native, Black or African American (including African and Caribbean), and Hispanic or Latino. Demographic groups who were not considered underrepresented in STEM included students who identified as White, Asian (including subcontinent and Philippines), and of Middle Eastern descent. Participants also reported whether either parent attended any college (continued-generation college students, 71%) or neither parent attended any college (first-generation college students, 29%). Moreover, student participants ranged in academic year (4% first-year students, 34% second-year students, 31% third-year students, 29% fourth-year students, and 2% students who already had a bachelor’s degree). For our study, we combined first- and second-year students (38% of participants) and third-year students and beyond (62%).

### Measure of research experience and dosage

Pre-course surveys asked participants to report any research-associated experiences prior to the Fly-CURE. Refer to pre-survey question 7 (Appendix 2) for the specific experiences listed. Students who chose “yes” to any of these experiences were considered as having prior research exposure, while those who did not choose “yes” to any of these questions were considered as not having prior research exposure.

### Fly-CURE outcome measures

Survey items for assessing research self-efficacy and sense of belonging were adapted from items used in the evaluation of the National Institutes of Health’s Building Infrastructure Leading to Diversity (BUILD) initiative. This retrospective pre-/post-survey method of measuring outcomes is commonly used when there is a possibility that students’ understanding of the constructs, such as what a research-intensive science laboratory course is, changes as a result of participating in the course and eliminates the possibility of a response shift bias in the data (38). For each evaluated outcome, students self-reported their pre- and post-course confidence or agreement with specific matrices using a 1-5 Likert scale.

#### Research self-efficacy

Pre- and post-course surveys asked students to report their perceived abilities and confidence for eight statements (Appendix 2, pre-survey question 8 and post-survey question 4). The scores from all eight questions were added together, resulting in a scale ranging from 8 to 40. Psychometric analysis of the pre- and post-course survey data revealed that this scale had a coefficient alpha of 0.918 for the pre-survey and 0.975 for the post-course survey, indicating these items measure the same construct.

#### Sense of belonging in science

Pre- and post-course surveys asked students to report their perceived agreement with four statements (Appendix 2, pre-survey question 9 and post-survey question 5). To determine scale scores, the results from all four questions were added together, resulting in a scale of 4 to 20. Psychometric analysis revealed that this scale had coefficient alphas of 0.863 and 0.935 for the pre- and post-course surveys, respectively.

#### Intent to pursue additional research opportunities

Post-course surveys asked participants to report their perceived intentions before and after taking the course. Students reported their likelihood to do each of the following: (i) enroll in another research-intensive science laboratory course; (ii) pursue or continue independent research in a science laboratory; (iii) pursue a career as a scientist (Appendix 2, post-survey questions 1-3). The scores from all three questions were analyzed separately or added together on a scale of 3 to 15. Psychometric analysis showed that this scale had a coefficient alpha of 0.861 for the pre-survey and 0.789 for the post-course survey.

### Statistical analyses

Independent groups and paired t-tests were used to assess the statistical significance of differences in the means within the same students from pre- to post-course (paired t-tests) and between different groups of students (independent groups t-tests). Levene’s Test for Equality of Variances was used to test for homogeneity of variance.

The mean scores for the three outcome scales were calculated in two ways: the scale score means and the gain score mean. Two scale score means are calculated for each outcome, a pre- and a post-course scale score mean, representing the average of student scale scores. The scale score mean may underestimate change because some students may have rated themselves the highest possible score on the pre-course survey. If they also rate themselves the highest possible score on the post-course survey, the difference between the pre- and post-course scores is zero. These students may have rated themselves even higher on the post-course survey, but the maximum possible score presented a ceiling for them. Thus, the scale score mean includes these zeros and deflates the mean score for the group. To account for this, a second mean score was calculated using the normalized gain score. The gain score removes students with the highest possible pre-course score from the analysis and examines the degree of change among students who *could* change because they did not reach the ceiling score on the pre-course survey (39). The equation used to calculate the normalized gain score is: *Normalized Gain = (Post-score - Pre-score)/(Maximum possible score - Pre-score)*. The data presented herein include both the scale score mean and the mean gain scores for all statistical comparisons.

## RESULTS

### The Fly-CURE focuses on the genomic mapping and phenotypic characterization of EMS-induced mutant lines involved in *Drosophila* eye development

At the beginning of each semester, all required *Drosophila* stocks were shipped to participating institutions. *Drosophila* mutant stocks contain previously generated EMS-induced mutations on the right arm of chromosome 2 (2R) (25). These mutations were previously identified based on homozygous recessive lethality and a growth-associated phenotype in the *Drosophila* eye when cell death is also blocked, but the genomic locus of the mutations is unknown (26, 27, 29–35). The identified mutants serve as the basis for phenotypic eye characterization, complementation mapping, and molecular analysis modules of the Fly-CURE (Figure 1A,B).

The Fly-CURE is a lab research project that includes both an initial “Discovery Phase” and a subsequent “Inquiry Phase” (Figure 1A). An initial pre-survey (Appendix 2) is first completed by all participating students to gather information about general student demographics, prior research experience, research self-efficacy, and sense of belonging in science. Students then typically complete an initial “Discovery Phase” of the project to characterize the eye tissue growth phenotype caused by the EMS-induced mutation and use complementation mapping of the lethal phenotype with a series of defined chromosomal deletions (40) to identify the genomic locus where the mutation responsible for the observed phenotype may be found. All recessive lethal EMS-induced mutations being investigated, as well as the chromosomal deficiencies used for complementation mapping, are maintained as heterozygotes using a second chromosome balancer causing curly wings (a dominant phenotypic marker; Figure 1A). Therefore, for crosses between the *Drosophila* mutant stock and stocks containing chromosomal deficiencies along 2R, students use stereomicroscopes to easily score for the presence (complementation) or absence (failure to complement) of straight-winged flies (those carrying the mutation and deficiency) among the progeny. Since the chromosomal deletions used in the first round of complementation mapping are relatively large and often lack several dozen to hundreds of genes (40), a second round of complementation tests with smaller deletions and/or chosen null alleles of individual genes within the specific genomic region identified in the first round of complementation mapping can be utilized to identify a smaller region where the mutation might be located. Once non-complementing deficiencies are identified, this concludes the “Discovery Phase” of the CURE.

During the “Inquiry Phase”, students develop hypotheses about candidate genes within the genomic region that fails to complement lethality of the mutation. Student-derived hypotheses usually focus on why mutations within a specific gene might lead to the observed eye tissue phenotype or recessive lethality. Typically, students choose genes that have been previously annotated as being involved in cellular growth control, apoptosis, the cell cycle, or similar processes. In some cases, the EMS mutation fails to complement a mutant allele of a specific gene by the second round of crosses (29, 32, 34, 35), allowing students to focus their hypothesis generation and subsequent molecular analyses on a single gene. Students then isolate genomic DNA from the mutant and control fly stocks, design PCR primers, and amplify a small (500-1000 nucleotide) region of their chosen gene. The sequence of the amplified region from both the mutant and control stocks is then determined by Sanger sequencing to identify possible differences between the heterozygous mutant stock and the wild-type control. Then, students use bioinformatics approaches to understand protein structure and/or evolutionary conservation of the candidate gene and often present their findings to the rest of the class. Finally, students analyze, summarize, and connect the data acquired. Different pedagogical assessments are used across the consortium, including formal lab reports, poster presentations, and micropublication-style manuscripts. At the end of the semester, a post-survey was completed to assess whether the semester-long Fly-CURE impacted students’ sense of belonging within the scientific community, feelings of self-efficacy in research, and motivation to pursue other future research experiences or STEM careers.

### Fly-CURE is a modular research experience that can be implemented in a variety of laboratory classes

The modular nature of Fly-CURE allows for components to be organized or omitted to meet the learning objectives and scheduling variability of different courses (Figure 1B). For example, most courses that have implemented the Fly-CURE have been upper-level genetics classes that also contain a laboratory component (Figure 1C, n=9). These combined lecture and lab courses, along with stand-alone genetics laboratory courses that lack a separate lecture component (n=4), typically utilize all modules of the Fly-CURE (Figure 1B, version 1). However, the Fly-CURE has also been implemented in Introductory Biology (n=1), a sophomore-level Molecular Biology course (n=1), and Anatomy and Physiology (n=1). In these non-genetics-centered classes, other variations of the Fly-CURE have been implemented that lack one or more of the modules contained in Fly-CURE version 1 (Figure 1B). Thus, while Fly-CURE has been mostly implemented in genetics courses, its adaptability and student-focused nature have allowed a wide variety of courses to participate in this course-embedded research experience.

While the modularity and adaptability of the Fly-CURE have allowed for its implementation in a variety of courses, we also wanted to assess whether faculty using this CURE could do so successfully without prior research experience with *Drosophila*. We surveyed faculty who had implemented the Fly-CURE and found that only slightly more than half (53%, n=8), had previously trained as a graduate student or postdoctoral fellow in a research lab where *Drosophila melanogaster* was utilized as a genetic model organism (Figure 1D). Together, these data suggest that Fly-CURE can be widely implemented in a variety of courses and that extensive prior training or experience in a *Drosophila* research lab by the faculty instructor is not a requisite for Fly-CURE implementation.

### The Fly-CURE provides research experiences at a range of institutions and for a broad spectrum of student participants

One motivation for the development of the Fly-CURE was to establish opportunities for collaboration between faculty and students at different institutions. Faculty were recruited to participate in Fly-CURE through a variety of methods, including discussions at conferences, social media, and word-of-mouth. The cohort of faculty collaborating on the Fly-CURE spanned several types of institutions (Figure 2A). Approximately equal numbers of faculty at Primarily Undergraduate Institutions (PUIs, n=6) and non-R2 graduate degree-granting institutions (n=5) have implemented the Fly-CURE into at least one course. In addition, the Fly-CURE has been implemented at R2 institutions (n=3) and at a community college (n=1), where undergraduate research experiences are typically limited due to a variety of factors including teaching load and institutional resources (3, 41, 42). Approximately 20% of institutions where the Fly-CURE has been taught over the last three years are also classified as Minority Serving Institutions (MSIs) (Figure 2B). Regular virtual meetings between participating faculty serve to foster collaboration between classes characterizing the same *Drosophila* mutation and have also culminated in eight collaborative micropublications consisting entirely of student-generated data (27, 29–35).

Among all students who have participated in the Fly-CURE, 27% self-identify as belonging to a demographic group underrepresented in STEM (Figure 2C) and 29% of students are first-generation college students (Figure 2D). In addition, only slightly more than half (52.5%) of students had any research exposure before the Fly-CURE (Figure 2E). Of the students who previously participated in a research experience, most had participated in a course-based research experience (Figure 2F), while only 26% of students had participated in a mentored apprenticeship-style research experience. Given the significant positive impacts that research experiences have on undergraduate STEM majors (43) and the dearth of mentored research experiences typically available to many undergraduate students, these data suggest that CUREs provide an important alternative to traditional apprentice-style research positions. While first-year undergraduate research experiences have been shown to be particularly important for the retention of STEM majors (44), the correlation between the number of research experiences a student participates in and student outcomes has been less well-studied. In particular, course-embedded research experiences like the Fly-CURE provide an additional “dose” of research to a large number of students, and in so doing, further promote student self-efficacy in research, sense of belonging in the scientific community, and pursuit of STEM careers.

### Impact of the Fly-CURE on student self-efficacy in research

To evaluate the impact of the Fly-CURE experience on students’ research self-efficacy, sense of belonging in science, and student interest in pursuing additional research experiences, pre- and/or post-course surveys were used to ask students about their confidence or level of agreement with multiple statements focused on these areas. Likert scale responses for questions focused on each metric were tallied to generate scale scores. Lower scale scores represent less confidence or agreement with associated statements, while higher scale scores represent students who reported more confidence or agreement with included statements.

As a first measurement of Fly-CURE effectiveness, we analyzed students’ sense of research self-efficacy. Students ranked their confidence in response to eight statements pertaining to this metric on pre- and post-course surveys (see Methods and Appendix 2). Students reported increased self-efficacy in research from pre-course to post-course, shown as an increase in scale score means (Figure 3A) and as a mean gain score (Figure 3B). We were also interested in whether the Fly-CURE closed gaps in research self-efficacy for specific student subgroups that are underrepresented in STEM, thereby providing a path to increased diversity in STEM. Interestingly, female students reported lower confidence in research pre-course (28.0 for females, 29.2 for males) and surpassed males in reported self-efficacy post-course (31.5 for females, 31.0 for males) (Figure 3C), resulting in a gain in research self-efficacy for both male and female students (Figure 3D). Although all student subgroups reported significant gains in their self-efficacy in research post-course, there were no statistically significant differences in the degree of reported gains in research self-efficacy between students in the evaluated subgroups, including race and ethnicity (Figure 3E, Supplemental Figure 1A,B), education background of parents (Figure 3E, Supplemental Figure 1C,D), and academic year (Supplemental Figure 1E,F).

**Figure 3.**
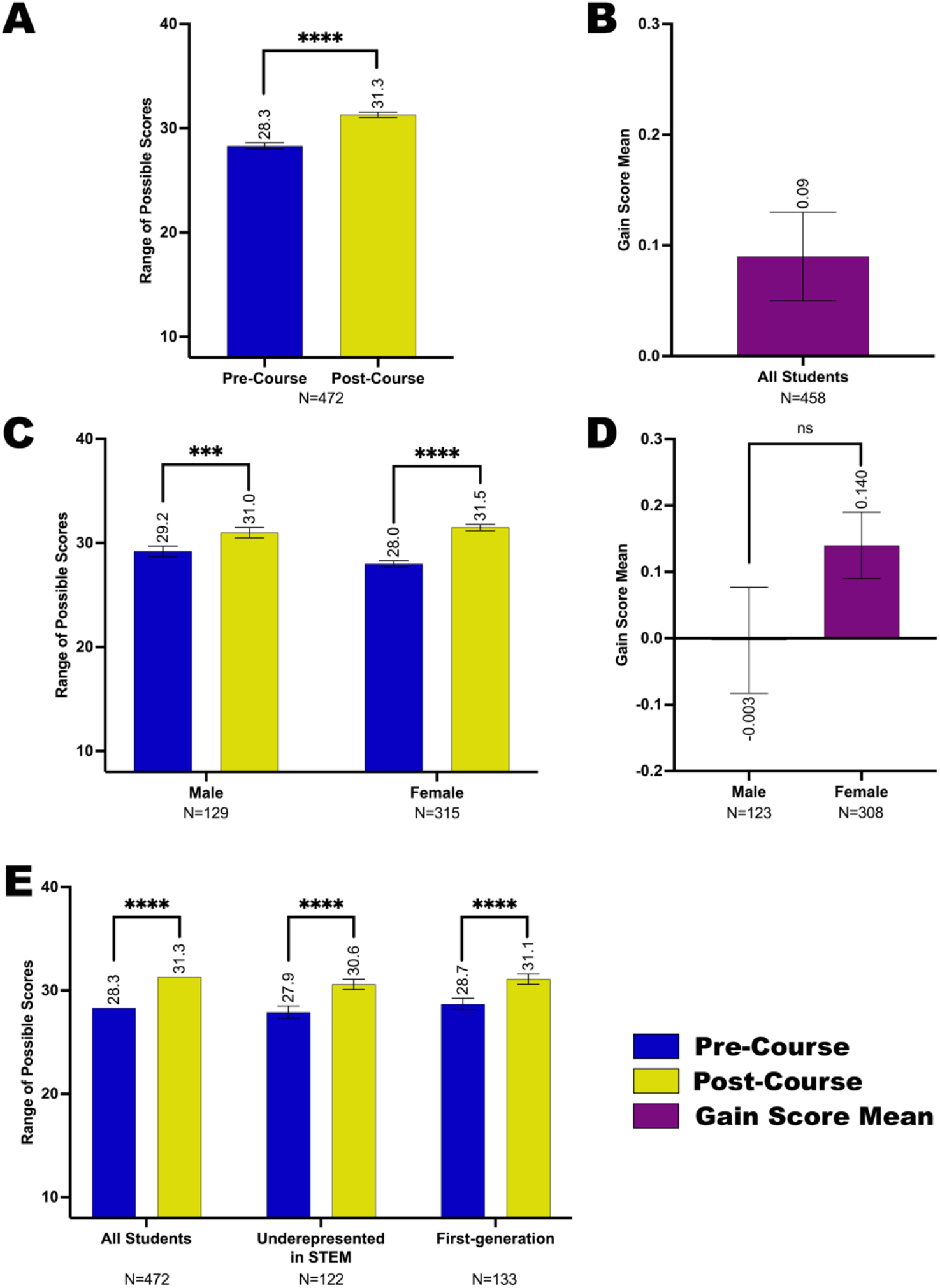
Self-efficacy in scientific research of student subgroups before and after completing the Fly-CURE. Through pre- and post-course surveys, students reported their efficacy in specific skills associated with scientific research before and after participating in the Fly-CURE. The survey rating scales for eight questions were combined, resulting in a total possible scale score of 40 (*y*-axis) per student. The mean self-efficacy pre-course (blue) and post-course (yellow) are shown for all participants (A) and in participant subgroups (C,E). (A-B) Self-efficacy scale score mean (A) and gain score mean (B) for all Fly-CURE participants. (C-D) Self-efficacy scale score mean (C) and gain score mean (D) for male and female participants. (E) Comparison of self-efficacy means pre- and post-course in all students, minority students underrepresented in STEM, and first-generation college students. Error bars represent ± standard error of the mean (± SEM); ns, not significant, *P* > 0.05; \*\*\*\**P* ≤ 0.0001.

### Impact of the Fly-CURE on student sense of belonging in the scientific community

Pre- and post-course surveys were also used to evaluate the effectiveness of the Fly-CURE in increasing student sense of belonging in science by asking students to rate their level of agreement with four statements (see Methods and Appendix 2). Pre- and post-course sense of belonging scales were generated by adding each student’s ratings on the four items.

Similar to their reported gains in research self-efficacy, students reported an increased sense of belonging in the scientific community post-course compared to pre-course. This is shown as scale score means (Figure 4A) and as a mean gain score (Figure 4B). We also compared student subgroups in several demographic categories and found that although all student subgroups reported gains in their feelings of belonging in science post-course, there were no statistically significant differences in the degree of reported gains between subgroups in each evaluated category, including gender (Figure 4C,D), race and ethnicity (Figure 4E, Supplemental Figure 2A-B), education background of parents (Figure 4E, Supplemental Figure 2C,D), and academic year (Supplemental Figure 2E,F). These data suggest that students from underrepresented backgrounds participating in Fly-CURE make similar gains as their peers. It is worth noting that similar to research self-efficacy, female participants reported a lower sense of belonging in science pre-course (12.2) compared to males (13.1), but yet reached a score similar to males post-course (13.8 for females, 14.0 for males) (Figure 4C). This suggests that the Fly-CURE experience allows female students to increase their perceived sense of belonging in science, thereby narrowing the gender gap in STEM.

**Figure 4.**
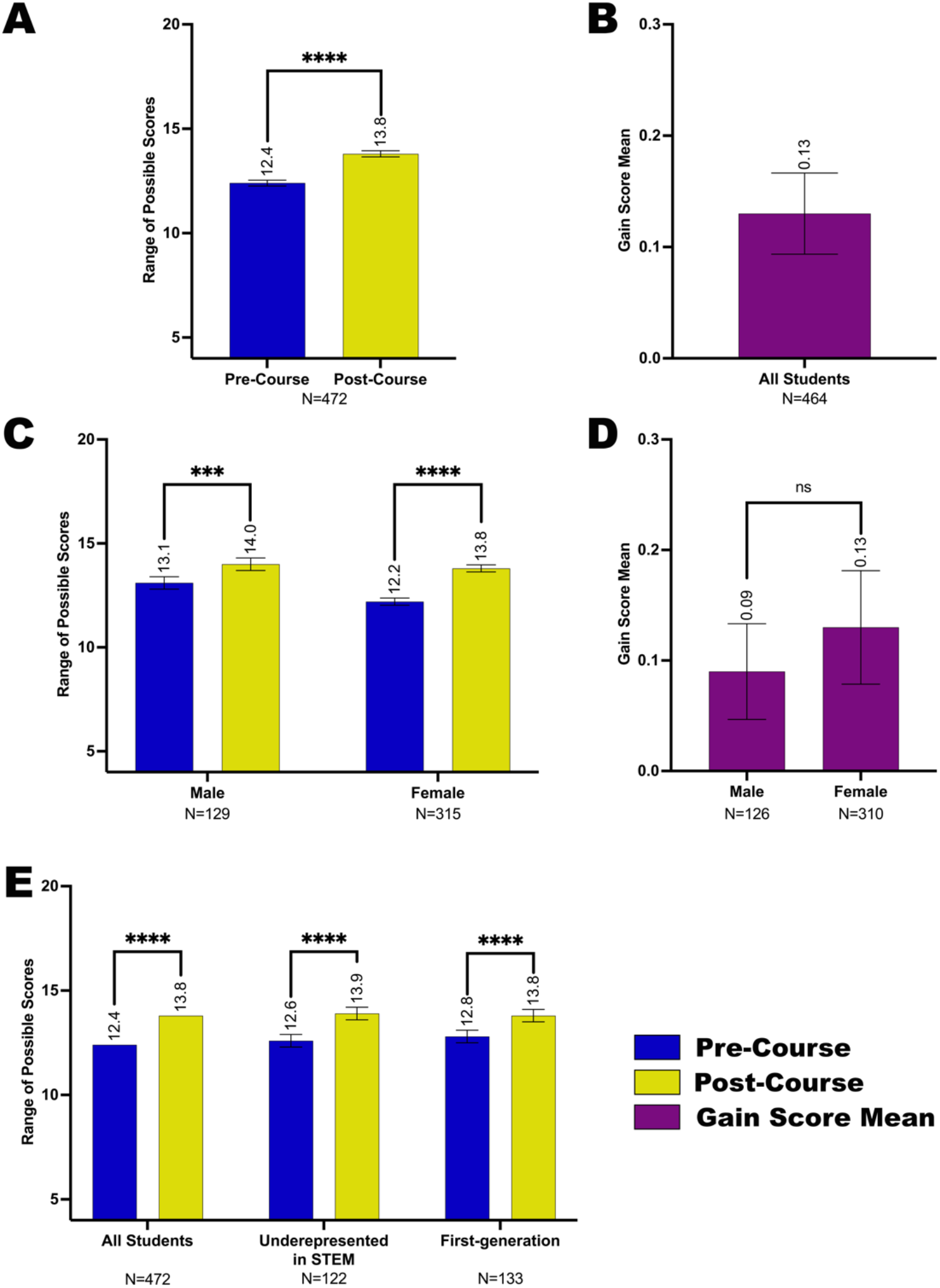
Sense of belonging in the scientific community for student subgroups before and after completing the Fly-CURE. Through pre- and post-course surveys, students reported their sense of belonging in the scientific community before and after participating in the Fly-CURE. The survey rating scales for four questions were combined, resulting in a total possible scale score of 20 (*y*-axis) per student. The mean scale score for sense of belonging pre-course (blue) and post-course (yellow) are shown for all participants (A) and in participant subgroups (C,E). (A-B) Sense of belonging scale score mean (A) and gain score mean (B) for all Fly-CURE participants. (C-D) Sense of belonging scale score mean (C) and gain score mean (D) for male and female students. (E) Comparison of reported scale score means for sense of belonging for all participants, minority students underrepresented in STEM, and first-generation college students. Error bars, ±SEM; ns, not significant, *P* > 0.05; \*\*\**P* ≤ 0.001; \*\*\*\**P* ≤ 0.0001.

### Impact of the Fly-CURE on student intention to pursue additional research opportunities

To evaluate the effectiveness of the Fly-CURE in increasing student intention to pursue additional research-associated experiences, post-course surveys asked participants to rate their perceived likelihood to seek out additional research opportunities before and after taking the course for three questions (see Methods and Appendix 2). Much like the reported gains in research self-efficacy and sense of belonging in science, students also reported a perceived increase in their intention to pursue additional research experiences after completing the Fly-CURE. This can be observed as scale score means (Figure 5A), as a mean for each type of experience evaluated (Figure 5B), and as a mean gain score for each type of experience (Figure 5C). It is worth noting that all student subgroups analyzed tend to start at a similar level of perceived intent to pursue the experiences proposed before the course and have a similar level of intent after the course (Supplemental Figure 3). Altogether, these data highlight the positive impact that the Fly-CURE has on encouraging confidence, belonging, and persistence in science for students who participate in a CURE during their undergraduate education.

**Figure 5.**
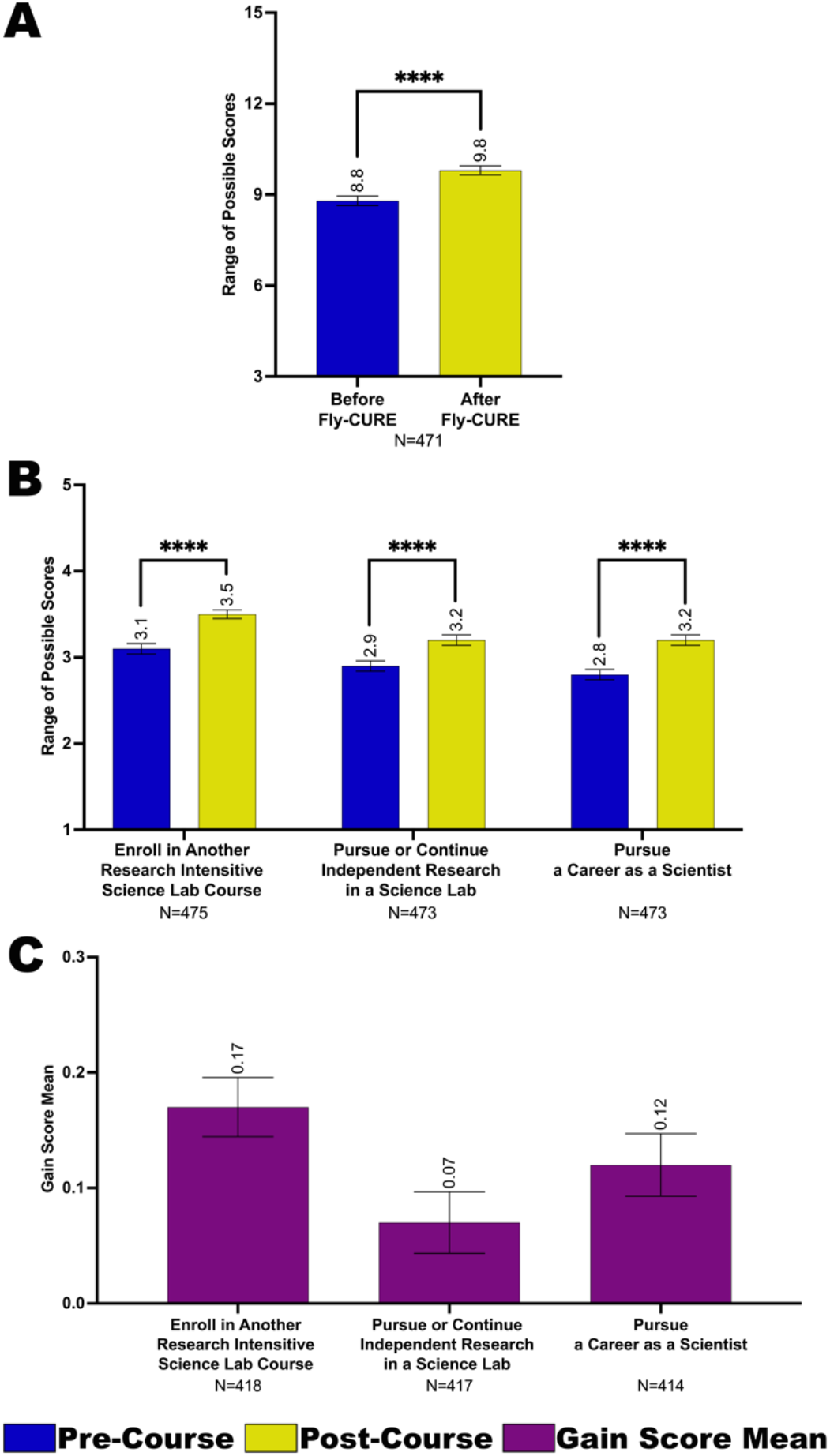
Student intent to seek additional research experiences before and after completing the Fly-CURE. Students reported their perceived interest in pursuing additional research–associated experiences before and after completing the Fly-CURE. The survey rating scales for three questions were combined, resulting in a maximum scale score of 15 (*y*-axis) per student. Students were asked to evaluate their perceived interest before and after the CURE in the categories listed in (B and C). (A-B) Scale score means for interest in seeking additional research experiences before (blue) compared to after (yellow) the Fly-CURE for all participants. (A) Scale score means across all categories. (B) Scale score means for individual categories evaluating student intent to seek additional research opportunities. (C) Gain score means comparing students’ interest in pursuing additional research experiences before and after the Fly-CURE for each category evaluated. Error bars, ±SEM; \**P* ≤ 0.05; \*\*\*\**P* ≤ 0.0001.

### Impact of the Fly-CURE on students with and without previous research experiences

While much of our data support previously reported impacts that CUREs have on student gains (42, 48), thereby highlighting the effectiveness of the Fly-CURE experience for students, we were also interested in evaluating the impacts of the Fly-CURE on students with or without research experience prior to taking a Fly-CURE course. In a pre-course survey, students were asked which specific research experiences, if any, they had prior to beginning the Fly-CURE project (see Methods and Appendix 2). Approximately 53% of students reported having had research experience of some kind before starting the Fly-CURE (Figure 2E).

Students with and without prior research experience reported gains in self-efficacy in research (Figure 6A) and sense of belonging in the scientific community (Figure 6C) after completing the Fly-CURE. Interestingly, however, students without prior research experience reported a greater gain in research self-efficacy after the Fly-CURE, suggesting that the Fly-CURE serves as a valuable research experience for those students and makes strides in increasing their confidence in conducting research (Figure 6B). On the contrary, students with and without prior research experience did not exhibit differential gains in their sense of belonging to the scientific community (Figure 6D). It is important to note, however, that the mean sense of belonging score for students without prior research experience post-course (13.3) surpassed the pre-course score for students with prior research experience (13.0) (Figure 6C). This indicates that the research experience component of the Fly-CURE increases students’ sense of belonging in science from baseline and suggests that there may be a dose-dependent relationship between the number of research experiences a student has and students’ sense of belonging in the scientific community.

**Figure 6.**
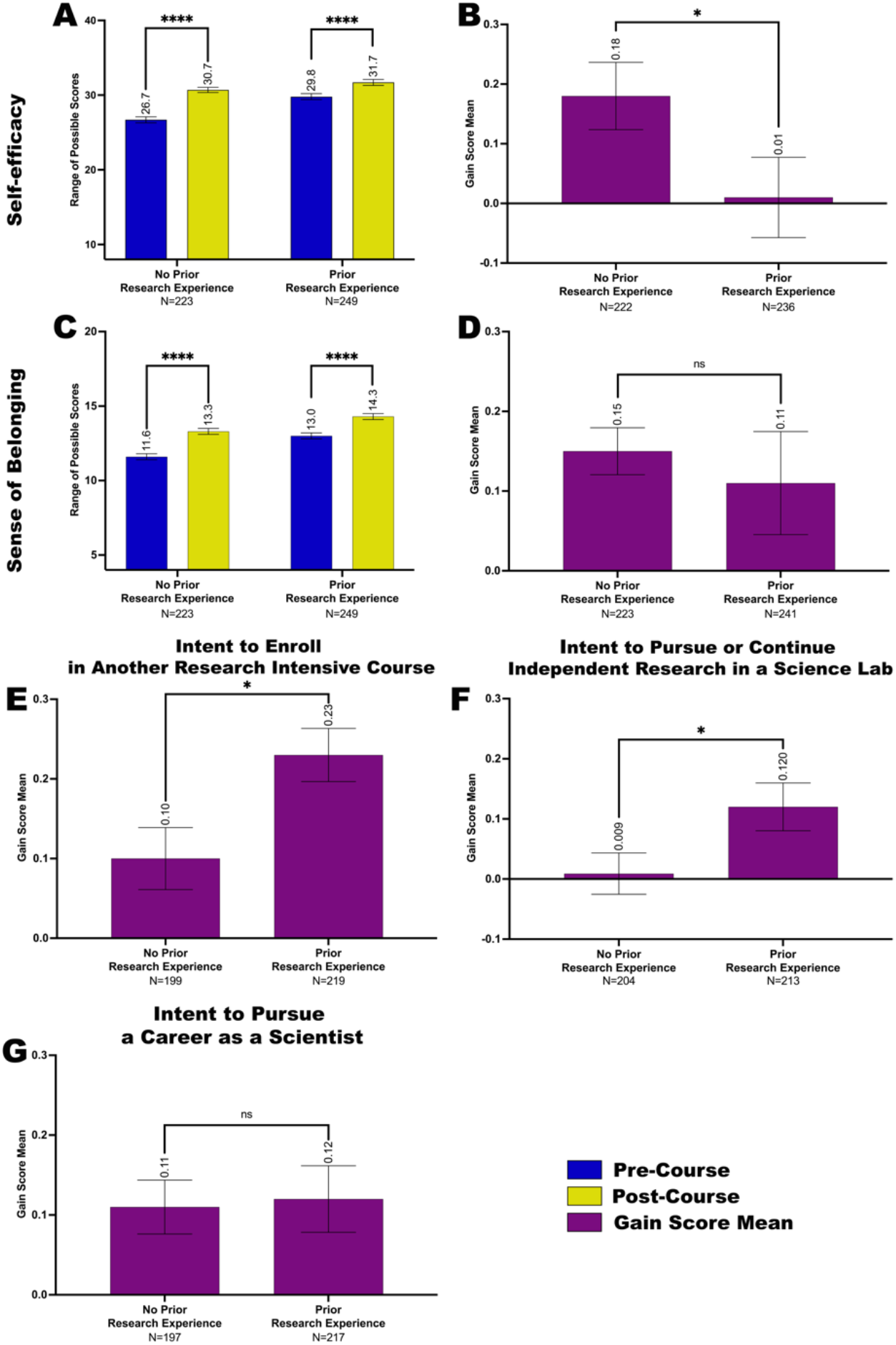
Impacts of self-efficacy in scientific research, sense of belonging in the scientific community, and intent to seek additional research experiences in students with and without research experience prior to the Fly-CURE. Through pre- and post-course surveys, students reported their self-efficacy in scientific research (A-B), sense of belonging in the scientific community (C-D), interest in pursuing additional research-associated experiences (E-G), and whether they had research experience prior to the course. (A,C) Scale score mean for research self-efficacy (A) and sense of belonging in science (C) before (blue) and after (yellow) the Fly-CURE for participants with and without prior research experience. (B,D) Gain score mean for self-efficacy (B) and sense of belonging (D) for Fly-CURE participants with and without prior research experience. (A) For research self-efficacy, the survey rating scales for eight questions were combined, resulting in a maximum score of 40 (*y*-axis). (C) For sense of belonging in science, the survey rating scales for four questions were summed, resulting in a combined score of 20 (*y*-axis). (E-G) Gain score means for students’ perceived interest to enroll in another research-intensive science laboratory course (E), pursue or continue independent research in a research laboratory (F), and pursue a career as a scientist (G) before and after taking the Fly-CURE. Error bars, ±SEM; ns, not significant, *P* > 0.05; \**P* ≤ 0.05; \*\*\*\**P* ≤ 0.0001.

Next, we evaluated whether the Fly-CURE had differing impacts on students’ intention to pursue additional research opportunities depending on whether students entered the CURE with or without prior research experience. In particular, we questioned whether participating in at least one research experience before the Fly-CURE resulted in a greater increase in students’ intent to seek out future research experiences compared to those without prior research experience. We found that although interest in gaining additional future experiences increased for all students (Figure 5A, Supplemental Figure 3), differential outcomes were observed, depending on the type of experience and whether students had research experience before taking the Fly-CURE course (Figure 6E-G). When students were asked whether they were interested in enrolling in another research-intensive laboratory course such as a CURE, students with prior research experience exhibited a greater gain in intent to pursue this experience, as shown by the gain score mean post-Fly-CURE compared to pre-Fly-CURE (*P* ≤ 0.05) (Figure 6E; scale score mean data in Supplemental Figure 4A). Similarly, students with prior research experience reported a greater gain in their intent to pursue or continue independent research in a scientific research laboratory than students without prior research experience (*P* ≤ 0.05) (Figure 6F; Supplemental Figure 4B). These data support the hypothesis that increased dosage in research experiences positively correlates with increases in student interest to persist in research. The only category in which students with and without prior research experience did not show differential outcomes was for intention to pursue a career as a scientist (Figure 6G; Supplemental Figure 4C). Regardless of the extent of prior research experience, students reported a very similar increase in intent to become scientists post-Fly-CURE as they did before Fly-CURE, suggesting that additional exposure to research does not significantly increase, beyond the initial positive impact, students’ interest to pursue a career in STEM.

Altogether, our data show that all students, regardless of demographic profile and previous exposure to research, show an increase in research self-efficacy, sense of belonging in science, and interest in pursuing additional research experiences after taking a Fly-CURE course. In addition, students without prior research experience show a statistically significant gain in self-efficacy compared with students with prior research experience; while students with prior experience in research show a statistically significant gain in interest to seek out additional research opportunities, but no significant increase in intent to pursue a career as a scientist.

## DISCUSSION

The Fly-CURE is a versatile authentic research experience that can be implemented in a modular fashion across course and/or institution types, and without requiring prior experience with *Drosophila melanogaster* (Figure 1B-D and Figure 2A,B). Thus, the Fly-CURE consortium is a large and diverse sample for measuring the impact of course-embedded research on student attitudes regarding self-efficacy in research, sense of belonging in science, intent to pursue additional research experiences, and the impact of previous research experiences (dosage) on these metrics. Prior studies have suggested that increased time spent on a task and research dosage positively impact student outcomes and persistence in STEM (45, 46). However, it has been suggested that persisting in science may require “a commitment of 10 or more hours per week over two or more semesters of faculty-mentored research” (3, 45). Therefore, we investigated the relationship between research exposure and its impacts on students’ retention, belonging, and confidence in STEM.

Overall, gains were reported by Fly-CURE students for scientific self-efficacy and sense of belonging, as well as for their intent to persist in STEM. Our analysis shows that all participating students, including groups considered underrepresented in STEM, females, and first-generation college students, reported increased confidence in research-associated skills (Figure 3 and Supplemental Figure 1), sense of belonging in science (Figure 4 and Supplemental Figure 2), and interest in pursuing additional research experiences (Figure 5 and Supplemental Figure 3) after the Fly-CURE. These are gains previously reported by others and our data supports the growing notion that CUREs are inclusive and have a positive impact on undergraduate STEM education (10, 11, 13–17).

Further, the fact that Fly-CURE is successfully implemented by faculty at a wide range of institutions (e.g., PUI, CC, MSI, and R2), a variety of courses, and by faculty without prior experience with *Drosophila* demonstrates the adaptable nature of the Fly-CURE. This also exemplifies the effectiveness of the Fly-CURE consortium for providing authentic research experiences for an increased number of STEM students. Traditional apprentice-based research experiences are often limited in availability, budget, and/or capacity, rendering the need for course-based experiences. However, one of the barriers to starting a CURE is having a project that is sustainable and feasible within the confines of an undergraduate curriculum. Additional barriers to CURE implementation exist for some institutional types such as community colleges (47). Nevertheless, community college students have comparable knowledge and perceived outcomes gains as non-community college counterparts when engaging in centrally supported CUREs, demonstrating the need for these research experiences to be accessible to all students (41, 42). The versatility associated with the modular nature of experiments in the Fly-CURE, as well as the diverse range of institutions at which the Fly-CURE has been implemented successfully, highlight its value for both students and curricula.

While other research endeavors have looked at dosage in terms of how much time a researcher spends on a single project (45, 46), we were able to investigate whether a separate previous research experience had an impact on changes in attitude resulting from the Fly-CURE (Figure 6 and Supplemental Figure 4). There were two lessons that emerged from our findings that could impact how undergraduate STEM departments incorporate research into their curriculum. The first is that students with no self-reported previous research experience demonstrated gains in both research self-efficacy and sense of belonging after a single semester of research (Figure 6A-D). Perhaps not surprisingly, these students reported a more significant gain in research self-efficacy than their classmates who had previous research experience (Figure 6A,B). This may be one of the most promising aspects of the Fly-CURE as a pedagogy to broaden participation in institutions where research opportunities are especially limited, such as two-year institutions with the most diverse student populations. Additionally, students with prior research experience reported a more significant gain in their intent to enroll in another research-intensive course and pursue independent research in a science lab (Figure 6E,F), highlighting a correlation between increased dosage and interest to persist in research. However, both students with and without prior research experience showed similar gains in their intent to pursue a career as a scientist (Figure 6G), suggesting that career plans might be less subjective to research exposure dosage. It is worth noting that the future career plans for many Fly-CURE participants might be in STEM-related careers, such as health professions, but not necessarily in laboratory research. Thereby, we predict that most respondents perceived a “career as a scientist” as a bench or field scientist, rather than a health-centered career. In the future, it would be enlightening to offer more specific career avenues to better appreciate the impact of the Fly-CURE on participants’ career interests.

Overall, these data show that participation in the Fly-CURE, as a single research experience, increases these metrics, even if this CURE is the student’s first research experience. Second, those students who had previous research experience also had statistically significant gains after completing the Fly-CURE, suggesting that all students have room to grow for the metrics analyzed in the second (or beyond) research experience. From our data, we cannot conclude how many research experiences would saturate these reported gains; however, we think it is reasonable to hypothesize that additional research experiences would result in additional gains in these areas. Future studies should specifically evaluate the critical number of research experiences associated with these and other student outcomes. Nonetheless, our data support previous evidence on the impacts of CUREs, thereby further underlining the importance for undergraduate STEM departments to incorporate one (or more) research experiences into the standardized curriculum.

## Supporting information

Combined Appendices

## ACKNOWLEDGEMENTS

We thank all Fly-CURE students who completed surveys and contributed to this work, as well as Alexandra Peister at Morehouse College and Clare Kron, Elizabeth Taylor, Hemin Shah, and Sandesh Pandit at NIU. Thanks also to the Bloomington *Drosophila* Stock Center, especially Kevin Cook and Cale Whitworth, for generously providing stocks. Figure 1A was made using BioRender.com. The research was supported by NSF IUSE grant 2021146 to JDK, KLB, JS, and ADV-M and NIH ReBUILDetroit grants UL1GM118982, TL4GM118983, and RL5GM118981 to JDK, VLS, and MAH.

## FIGURE LEGENDS

**Figure S1. Reported self-efficacy in research for student subgroups before and after completing the Fly-CURE.** The mean self-efficacy pre-course (blue) and post-course (yellow) scale score means are shown for participant subgroups, as well as the gain score mean (purple) to compare differential gains in self-efficacy post-course compared to pre-course in student subgroups. (A-B) Self-efficacy scale score means (A) and gain score mean (B) for minority students underrepresented in STEM and students not considered underrepresented in STEM. (C-D) Self-efficacy scale score mean (C) and gain score mean (D) for first-generation and continued-generation college students. (E-F) Self-efficacy scale score mean (E) and gain score mean (F) for first- or second-year students compared to third-year students and above. Error bars, ±SEM; ns, not significant, *P* > 0.05; \*\*\*\**P* ≤ 0.0001.

**Figure S2. Reported sense of belonging in science for student subgroups before and after completing the Fly-CURE.** The scale score means for sense of belonging in the scientific community pre-course (blue) and post-course (yellow) are shown for participant subgroups, as well as the gain score mean (purple) to compare differential gains in sense of belonging post-course compared to pre-course in student subgroups. (A-B) Sense of belonging scale score mean (A) and gain score mean (B) for minority students underrepresented in STEM and students not considered underrepresented in STEM. (C-D) Sense of belonging in research scale score mean (C) and gain score mean (D) for first-generation and continued-generation college students. (E-F) Sense of belonging scale score mean (E) and gain score mean (F) for first- or second-year students compared to third-year students and above. Error bars, ±SEM; ns, not significant, *P* > 0.05; \*\*\**P* ≤ 0.001; \*\*\*\**P* ≤ 0.0001.

**Figure S3. Reported intent to seek additional research experiences for student subgroups before and after completing the Fly-CURE.** Comparison of students’ perceived interest before (blue) and after (yellow) the CURE to enroll in another research-intensive science laboratory course (A,D,G,J), pursue or continue independent research in a research laboratory (B,E,H,K), and pursue a career as a scientist (C,F,I,L). Scale score means for reported perceived interest in seeking additional research experiences before compared to after the Fly-CURE for male and female students (A-C), minority students underrepresented in STEM and students not considered underrepresented in STEM (D-F), first-generation and continued-generation college students (G-I), and first- or second-year students compared to third-year students and above (J-L). Error bars, ±SEM; ns, not significant, *P* > 0.05; \**P* ≤ 0.05; \*\**P* ≤ 0.01; \*\*\**P* ≤ 0.001; \*\*\*\**P* ≤ 0.0001.

**Figure S4. Reported intent to seek additional research experiences in students with and without research experience prior to the Fly-CURE.** Through post-surveys, students reported their perceived interest in pursuing additional research–associated experiences before and after completing the Fly-CURE. The survey rating scales ranged from one (not likely) to five (definitely) for the research experiences indicated. Scale score means of perceived student interest to enroll in another research-intensive science laboratory course (A), pursue or continue independent research in a research laboratory (B), and pursue a career as a scientist (C) before (blue) and after (yellow) taking the Fly-CURE for students who reported as having or not having research experience prior to the Fly-CURE. Error bars, ±SEM; \**P* ≤ 0.05; \*\*\*\**P* ≤ 0.0001.

## SUPPLEMENTAL MATERIALS

Appendix 1: Institutional and student demographics tables

Appendix 2: Pre- and post-surveys

Supplemental Figures 1-4

## REFERENCES

1. Woodin T, Carter VC, Fletcher L. 2010. Vision and Change in Biology Undergraduate Education, A Call for Action—Initial Responses. CBE—Life Sci Educ 9:71–73.

2. Vision and Change in Undergraduate Biology Education » About V&C: A Call to Action (2011). https://visionandchange.org/about-vc-a-call-to-action-2011/. Retrieved 29 November 2022.

3. Bangera G, Brownell SE. 2014. Course-Based Undergraduate Research Experiences Can Make Scientific Research More Inclusive. CBE—Life Sci Educ 13:602–606.

4. Elgin SCR, Hauser C, Holzen TM, Jones C, Kleinschmit A, Leatherman J. 2017. The GEP: Crowd-Sourcing Big Data Analysis with Undergraduates. Trends Genet 33:81–85.

5. Shaffer CD, Alvarez C, Bailey C, Barnard D, Bhalla S, Chandrasekaran C, Chandrasekaran V, Chung H-M, Dorer DR, Du C, Eckdahl TT, Poet JL, Frohlich D, Goodman AL, Gosser Y, Hauser C, Hoopes LLM, Johnson D, Jones CJ, Kaehler M, Kokan N, Kopp OR, Kuleck GA, McNeil G, Moss R, Myka JL, Nagengast A, Morris R, Overvoorde PJ, Shoop E, Parrish S, Reed K, Regisford EG, Revie D, Rosenwald AG, Saville K, Schroeder S, Shaw M, Skuse G, Smith C, Smith M, Spana EP, Spratt M, Stamm J, Thompson JS, Wawersik M, Wilson BA, Youngblom J, Leung W, Buhler J, Mardis ER, Lopatto D, Elgin SCR. 2010. The Genomics Education Partnership: Successful Integration of Research into Laboratory Classes at a Diverse Group of Undergraduate Institutions. CBE—Life Sci Educ 9:55–69.

6. Tootle T, Hoffmann D, Allen A, Spracklen A, Groen C, Kelpsch D. 2019. Research and Teaching: Mini-Course-Based Undergraduate Research Experience: Impact on Student Understanding of STEM Research and Interest in STEM Programs. J Coll Sci Teach 048.

7. Delventhal R, Steinhauer J. 2020. A course-based undergraduate research experience examining neurodegeneration in *Drosophila melanogaster* teaches students to think, communicate, and perform like scientists. PLOS ONE 15:e0230912.

8. Mills A, Jaganatha V, Cortez A, Guzman M, Burnette JM, Collin M, Lopez-Lopez B, Wessler SR, Van Norman JM, Nelson DC, Rasmussen CG. 2021. A Course-Based Undergraduate Research Experience in CRISPR-Cas9 Experimental Design to Support Reverse Genetic Studies in *Arabidopsis thaliana*. J Microbiol Biol Educ 22:e00155–21.

9. Murren CJ, Wolyniak MJ, Rutter MT, Bisner AM, Callahan HS, Strand AE, Corwin LA. 2019. Undergraduates Phenotyping *Arabidopsis* Knockouts in a Course-Based Undergraduate Research Experience: Exploring Plant Fitness and Vigor Using Quantitative Phenotyping Methods. J Microbiol Biol Educ 20:10.

10. Brownell SE, Hekmat-Scafe DS, Singla V, Chandler Seawell P, Conklin Imam JF, Eddy SL, Stearns T, Cyert MS. 2015. A High-Enrollment Course-Based Undergraduate Research Experience Improves Student Conceptions of Scientific Thinking and Ability to Interpret Data. CBE—Life Sci Educ 14:ar21, 1–14.

11. Hanauer DI, Graham MJ, SEA-PHAGES, Betancur L, Bobrownicki A, Cresawn SG, Garlena RA, Jacobs-Sera D, Kaufmann N, Pope WH, Russell DA, Jacobs WR, Sivanathan V, Asai DJ, Hatfull GF, Actis L, Adair T, Adams S, Alvey R, Anders K, Anderson WA, Antoniacci L, Ayuk M, Baliraine F, Balish M, Ball S, Barbazuk B, Barekzi N, Barrera A, Berkes C, Best A, Bhalla S, Blumer L, Bollivar D, Bonilla JA, Borges K, Bortz B, Breakwell D, Breitenberger C, Breton T, Brey C, Bricker JS, Briggs L, Broderick E, Brooks TD, Brown-Kennerly V, Buckholt M, Butela K, Byrum C, Cain D, Carson S, Caruso S, Caslake L, Chia C, Chung H-M, Clase K, Clement B, Conant S, Connors B, Coomans R, D’Angelo W, D’Elia T, Daniels CJ, Daniels L, Davis B, DeCourcy K, DeJong R, Delaney-Nguyen K, Delesalle V, Diaz A, Dickson L, Doty J, Doyle E, Dunbar D, Easterwood J, Eckardt M, Edgington N, Elgin S, Erb M, Erill I, Fast K, Fillman C, Findley A, Fisher E, Fleischacker C, Fogarty M, Frederick G, Frost V, Furbee E, Gainey M, Gallegos I, Gissendanner C, Golebiewska U, Grose J, Grubb S, Guild N, Gurney S, Hartzog G, Hatherill JR, Hauser C, Hendrickson H, Herren C, Hinz J, Ho E, Hope S, Hughes L, Hull A, Hutchison K, Isern S, Janssen G, Jarvik J, Johnson A, Jones N, Kagey J, Kart M, Katsanos J, Keener T, Kenna M, King R, King-Smith C, Kirkpatrick B, Klyczek K, Koch H, Koga A, Korey C, Krukonis G, Kurt B, Leadon S, LeBlanc-Straceski J, Lee J, Lee-Soety J, Lewis L, Limeri L, Little J, Llano M, Lopez J, MacLaren C, Makemson J, Martin S, Mavrodi D, McGuier N, McKinney A, McLean J, Merkhofer E, Michael S, Miller E, Mohan S, Molloy S, Monsen-Collar K, Monti D, Moyer A, Neitzel J, Nelson P, Newman R, Noordewier B, Olapade O, Ospina-Giraldo M, Page S, Paige-Anderson C, Pape-Zambito D, Park P, Parker J, Pedulla M, Peister A, Pfaffle P, Pirino G, Pizzorno M, Plymale R, Pogliano J, Pogliano K, Powell A, Poxleitner M, Preuss M, Reyna N, Rickus J, Rinehart C, Robinson C, Rodriguez-Lanetty M, Rosas-Acosta G, Ross J, Rowland N, Royer D, Rubin M, Sadana R, Saha M, Saha S, Sandel M, Sasek T, Saunders L, Saville K, Scherer A, Schildbach J, Schroeder S, Schwebach JR, Seegulam M, Segura-Totten M, Shaffer C, Shanks R, Sipprell A, Slowan-Pomeroy T, Smith K, Smith MA, Smith-Caldas M, Stamm J, Stockwell S, Stowe E, Stukey J, Sunnen CN, Tarbox B, Taylor S, Temple L, Timmerman M, Tobiason D, Tolsma S, Torres M, Twichell C, Valle-Rivera AM, Vazquez E, Villagomez J, Voshell S, Wallen J, Ward R, Ware V, Warner M, Washington J, Weir S, Wertz J, Westholm D, Weston-Hafer K, Westover K, Whitefleet-Smith J, Wiedemeier A, Wolyniak M, Yan W, Zegers GP, Zhang D, Zimmerman A. 2017. An inclusive Research Education Community (iREC): Impact of the SEA-PHAGES program on research outcomes and student learning. Proc Natl Acad Sci 114:13531–13536.

12. Duboue ER, Kowalko JE, Keene AC. 2022. Course-based undergraduate research experiences (CURES) as a pathway to diversify science. Evol Dev 24:127–130.

13. Jordan TC, Burnett SH, Carson S, Caruso SM, Clase K, DeJong RJ, Dennehy JJ, Denver DR, Dunbar D, Elgin SCR, Findley AM, Gissendanner CR, Golebiewska UP, Guild N, Hartzog GA, Grillo WH, Hollowell GP, Hughes LE, Johnson A, King RA, Lewis LO, Li W, Rosenzweig F, Rubin MR, Saha MS, Sandoz J, Shaffer CD, Taylor B, Temple L, Vazquez E, Ware VC, Barker LP, Bradley KW, Jacobs-Sera D, Pope WH, Russell DA, Cresawn SG, Lopatto D, Bailey CP, Hatfull GF. 2014. A Broadly Implementable Research Course in Phage Discovery and Genomics for First-Year Undergraduate Students. mBio 5:e01051–13.

14. Cooper KM, Knope ML, Munstermann MJ, Brownell SE. 2020. Students Who Analyze Their Own Data in a Course-Based Undergraduate Research Experience (CURE) Show Gains in Scientific Identity and Emotional Ownership of Research. J Microbiol Biol Educ 21:60.

15. Bhatt JM, Challa AK. 2018. First Year Course-Based Undergraduate Research Experience (CURE) Using the CRISPR/Cas9 Genome Engineering Technology in Zebrafish. J Microbiol Biol Educ 19:19.1.30.

16. Evans CJ, Olson JM, Mondal BC, Kandimalla P, Abbasi A, Abdusamad MM, Acosta O, Ainsworth JA, Akram HM, Albert RB, Alegria-Leal E, Alexander KY, Ayala AC, Balashova NS, Barber RM, Bassi H, Bennion SP, Beyder M, Bhatt KV, Bhoot C, Bradshaw AW, Brannigan TG, Cao B, Cashell YY, Chai T, Chan AW, Chan C, Chang I, Chang J, Chang MT, Chang PW, Chang S, Chari N, Chassiakos AJ, Chen IE, Chen VK, Chen Z, Cheng MR, Chiang M, Chiu V, Choi S, Chung JH, Contreras L, Corona E, Cruz CJ, Cruz RL, Dang JM, Dasari SP, De La Fuente JRO, Del Rio OMA, Dennis ER, Dertsakyan PS, Dey I, Distler RS, Dong Z, Dorman LC, Douglass MA, Ehresman AB, Fu IH, Fua A, Full SM, Ghaffari-Rafi A, Ghani AA, Giap B, Gill S, Gill ZS, Gills NJ, Godavarthi S, Golnazarian T, Goyal R, Gray R, Grunfeld AM, Gu KM, Gutierrez NC, Ha AN, Hamid I, Hanson A, Hao C, He C, He M, Hedtke JP, Hernandez YK, Hlaing H, Hobby FA, Hoi K, Hope AC, Hosseinian SM, Hsu A, Hsueh J, Hu E, Hu SS, Huang S, Huang W, Huynh M, Javier C, Jeon NE, Ji S, Johal J, John A, Johnson L, Kadakia S, Kakade N, Kamel S, Kaur R, Khatra JS, Kho JA, Kim C, Kim EJ-K, Kim HJ, Kim HW, Kim JH, Kim SA, Kim WK, Kit B, La C, Lai J, Lam V, Le NK, Lee CJ, Lee D, Lee DY, Lee J, Lee J, Lee J, Lee J-Y, Lee S, Lee TC, Lee V, Li AJ, Li J, Libro AM, Lien IC, Lim M, Lin JM, Liu CY, Liu SC, Louie I, Lu SW, Luo WY, Luu T, Madrigal JT, Mai Y, Miya DI, Mohammadi M, Mohanta S, Mokwena T, Montoya T, Mould DL, Murata MR, Muthaiya J, Naicker S, Neebe MR, Ngo A, Ngo DQ, Ngo JA, Nguyen AT, Nguyen HCX, Nguyen RH, Nguyen TTT, Nguyen VT, Nishida K, Oh S-K, Omi KM, Onglatco MC, Almazan GO, Paguntalan J, Panchal M, Pang S, Parikh HB, Patel PD, Patel TH, Petersen JE, Pham S, Phan-Everson TM, Pokhriyal M, Popovich DW, Quaal AT, Querubin K, Resendiz A, Riabkova N, Rong F, Salarkia S, Sama N, Sang E, Sanville DA, Schoen ER, Shen Z, Siangchin K, Sibal G, Sin G, Sjarif J, Smith CJ, Soeboer AN, Sosa C, Spitters D, Stender B, Su CC, Summapund J, Sun BJ, Sutanto C, Tan JS, Tan NL, Tangmatitam P, Trac CK, Tran C, Tran D, Tran D, Tran V, Truong PA, Tsai BL, Tsai P-H, Tsui CK, Uriu JK, Venkatesh S, Vo M, Vo N-T, Vo P, Voros TC, Wan Y, Wang E, Wang J, Wang MK, Wang Y, Wei S, Wilson MN, Wong D, Wu E, Xing H, Xu JP, Yaftaly S, Yan K, Yang E, Yang R, Yao T, Yeo P, Yip V, Yogi P, Young GC, Yung MM, Zai A, Zhang C, Zhang XX, Zhao Z, Zhou R, Zhou Z, Abutouk M, Aguirre B, Ao C, Baranoff A, Beniwal A, Cai Z, Chan R, Chien KC, Chaudhary U, Chin P, Chowdhury P, Dalie J, Du EY, Estrada A, Feng E, Ghaly M, Graf R, Hernandez E, Herrera K, Ho VW, Honeychurch K, Hou Y, Huang JM, Ishii M, James N, Jang G-E, Jin D, Juarez J, Kesaf AE, Khalsa SK, Kim H, Kovsky J, Kuang CL, Kumar S, Lam G, Lee C, Lee G, Li L, Lin J, Liu J, Ly J, Ma A, Markovic H, Medina C, Mungcal J, Naranbaatar B, Patel K, Petersen L, Phan A, Phung M, Priasti N, Ruano N, Salim T, Schnell K, Shah P, Shen J, Stutzman N, Sukhina A, Tian R, Vega-Loza A, Wang J, Wang J, Watanabe R, Wei B, Xie L, Ye J, Zhao J, Zimmerman J, Bracken C, Capili J, Char A, Chen M, Huang P, Ji S, Kim E, Kim K, Ko J, Laput SLG, Law S, Lee SK, Lee O, Lim D, Lin E, Marik K, Mytych J, O’Laughlin A, Pak J, Park C, Ryu R, Shinde A, Sosa M, Waite N, Williams M, Wong R, Woo J, Woo J, Yepuri V, Yim D, Huynh D, Wijiewarnasurya D, Shapiro C, Levis-Fitzgerald M, Jaworski L, Lopatto D, Clark IE, Johnson T, Banerjee U. 2021. A functional genomics screen identifying blood cell development genes in *Drosophila* by undergraduates participating in a course-based research experience. G3 Genes Genomes Genet 11:jkaa028, 1–23.

17. Olson JM, Evans CJ, Ngo KT, Kim HJ, Nguyen JD, Gurley KGH, Ta T, Patel V, Han L, Truong-N KT, Liang L, Chu MK, Lam H, Ahn HG, Banerjee AK, Choi IY, Kelley RG, Moridzadeh N, Khan AM, Khan O, Lee S, Johnson EB, Tigranyan A, Wang J, Gandhi AD, Padhiar MM, Calvopina JH, Sumra K, Ou K, Wu JC, Dickan JN, Ahmadi SM, Allen DN, Mai VT, Ansari S, Yeh G, Yoon E, Gon K, Yu JY, He J, Zaretsky JM, Lee NE, Kuoy E, Patananan AN, Sitz D, Tran P, Do M-T, Akhave SJ, Alvarez SD, Asem B, Asem N, Azarian NA, Babaesfahani A, Bahrami A, Bhamra M, Bhargava R, Bhatia R, Bhatia S, Bumacod N, Caine JJ, Caldwell TA, Calica NA, Calonico EM, Chan C, Chan HH-L, Chang A, Chang C, Chang D, Chang JS, Charania N, Chen JY, Chen K, Chen L, Chen Y, Cheung DJ, Cheung JJ, Chew JJ, Chew NB, Chien C-AT, Chin AM, Chin CJ, Cho Y, Chou MT, Chow K-HK, Chu C, Chu DM, Chu V, Chuang K, Chugh AS, Cubberly MR, Daniel MG, Datta S, Dhaliwal R, Dinh J, Dixit D, Dowling E, Feng M, From CM, Furukawa D, Gaddipati H, Gevorgyan L, Ghaznavi Z, Ghosh T, Gill J, Groves DJ, Gurara KK, Haghighi AR, Havard AL, Heyrani N, Hioe T, Hong K, Houman JJ, Howland M, Hsia EL, Hsueh J, Hu S, Huang AJ, Huynh JC, Huynh J, Iwuchukwu C, Jang MJ, Jiang AA, Kahlon S, Kao P-Y, Kaur M, Keehn MG, Kim EJ, Kim H, Kim MJ, Kim SJ, Kitich A, Kornberg RA, Kouzelos NG, Kuon J, Lau B, Lau RK, Law R, Le HD, Le R, Lee C, Lee C, Lee GE, Lee K, Lee MJ, Lee RV, Lee SHK, Lee SK, Lee S-LD, Lee YJ, Leong MJ, Li DM, Li H, Liang X, Lin E, Lin MM, Lin P, Lin T, Lu S, Luong SS, Ma JS, Ma L, Maghen JN, Mallam S, Mann S, Melehani JH, Miller RC, Mittal N, Moazez CM, Moon S, Moridzadeh R, Ngo K, Nguyen HH, Nguyen K, Nguyen TH, Nieh AW, Niu I, Oh S-K, Ong JR, Oyama RK, Park J, Park YA, Passmore KA, Patel A, Patel AA, Patel D, Patel T, Peterson KE, Pham AH, Pham SV, Phuphanich ME, Poria ND, Pourzia A, Ragland V, Ranat RD, Rice CM, Roh D, Rojhani S, Sadri L, Saguros A, Saifee Z, Sandhu M, Scruggs B, Scully LM, Shih V, Shin BA, Sholklapper T, Singh H, Singh S, Snyder SL, Sobotka KF, Song SH, Sukumar S, Sullivan HC, Sy M, Tan H, Taylor SK, Thaker SK, Thakore T, Tong GE, Tran JN, Tran J, Tran TD, Tran V, Trang CL, Trinh HG, Trinh P, Tseng H-CH, Uotani TT, Uraizee AV, Vu KKT, Vu KKT, Wadhwani K, Walia PK, Wang RS, Wang S, Wang SJ, Wiredja DD, Wong AL, Wu D, Xue X, Yanez G, Yang Y-H, Ye Z, Yee VW, Yeh C, Zhao Y, Zheng X, Ziegenbalg A, Alkali J, Azizkhanian I, Bhakta A, Berry L, Castillo R, Darwish S, Dickinson H, Dutta R, Ghosh RK, Guerin R, Hofman J, Iwamoto G, Kang S, Kim A, Kim B, Kim H, Kim K, Kim S, Ko J, Koenig M, LaRiviere A, Lee C, Lee J, Lung B, Mittelman M, Murata M, Park Y, Rothberg D, Sprung-Keyser B, Thaker K, Yip V, Picard P, Diep F, Villarasa N, Hartenstein V, Shapiro C, Levis-Fitzgerald M, Jaworski L, Loppato D, Clark IE, Banerjee U. 2019. Expression-Based Cell Lineage Analysis in *Drosophila* Through a Course-Based Research Experience for Early Undergraduates. G3 Genes Genomes Genet 9:3791–3800.

18. Rodenbusch SE, Hernandez PR, Simmons SL, Dolan EL. 2016. Early Engagement in Course-Based Research Increases Graduation Rates and Completion of Science, Engineering, and Mathematics Degrees. CBE Life Sci Educ 15:ar20.

19. Freeman EA, Theodosiou NA, Anderson WJ. 2020. From bench to board-side: Academic teaching careers. Dev Biol 459:43–48.

20. Shortlidge EE, Bangera G, Brownell SE. 2017. Each to Their Own CURE: Faculty Who Teach Course-Based Undergraduate Research Experiences Report Why You Too Should Teach a CURE. J Microbiol Biol Educ 18:18.2.29.

21. Shortlidge EE, Bangera G, Brownell SE. 2016. Faculty Perspectives on Developing and Teaching Course-Based Undergraduate Research Experiences. BioScience 66:54–62.

22. Lopatto D, Alvarez C, Barnard D, Chandrasekaran C, Chung H-M, Du C, Eckdahl T, Goodman AL, Hauser C, Jones CJ, Kopp OR, Kuleck GA, McNeil G, Morris R, Myka JL, Nagengast A, Overvoorde PJ, Poet JL, Reed K, Regisford G, Revie D, Rosenwald A, Saville K, Shaw M, Skuse GR, Smith C, Smith M, Spratt M, Stamm J, Thompson JS, Wilson BA, Witkowski C, Youngblom J, Leung W, Shaffer CD, Buhler J, Mardis E, Elgin SCR. 2008. Genomics Education Partnership. Science 322:684–685.

23. Hanauer DI, Graham MJ, Arnold RJ, Ayuk MA, Balish MF, Beyer AR, Butela KA, Byrum CA, Chia CP, Chung H-M, Clase KL, Conant S, Coomans RJ, D’Elia T, Diaz J, Diaz A, Doty JA, Edgington NP, Edwards DC, Eivazova E, Emmons CB, Fast KM, Fisher EJ, Fleischacker CL, Frederick GD, Freise AC, Gainey MD, Gissendanner CR, Golebiewska UP, Guild NA, Hendrickson HL, Herren CD, Hopson-Fernandes MS, Hughes LE, Jacobs-Sera D, Johnson AA, Kirkpatrick BL, Klyczek KK, Koga AP, Kotturi H, LeBlanc-Straceski J, Lee-Soety JY, Leonard JE, Mastropaolo MD, Merkhofer EC, Michael SF, Mitchell JC, Mohan S, Monti DL, Noutsos C, Nsa IY, Peters NT, Plymale R, Pollenz RS, Porter ML, Rinehart CA, Rosas-Acosta G, Ross JF, Rubin MR, Scherer AE, Schroeder SC, Shaffer CD, Sprenkle AB, Sunnen CN, Swerdlow SJ, Tobiason D, Tolsma SS, Tsourkas PK, Ward RE, Ware VC, Warner MH, Washington JM, Westover KM, White SJ, Whitefleet-Smith JL, Williams DC, Wolyniak MJ, Zeilstra-Ryalls JH, Asai DJ, Hatfull GF, Sivanathan V. 2022. Instructional Models for Course-Based Research Experience (CRE) Teaching. CBE—Life Sci Educ 21:ar8, 1–14.

24. Caruso JP, Israel N, Rowland K, Lovelace MJ, Saunders MJ. 2016. Citizen Science: The Small World Initiative Improved Lecture Grades and California Critical Thinking Skills Test Scores of Nonscience Major Students at Florida Atlantic University. J Microbiol Biol Educ 17:156–162.

25. Neufeld TP, Hariharan IK. 2002. Regulation of Growth and Cell Proliferation During Eye Development, p. 107–133. In Moses, K (ed.), Drosophila Eye Development. Springer Berlin Heidelberg, Berlin, Heidelberg.

26. Kagey JD, Brown JA, Moberg KH. 2012. Regulation of Yorkie activity in *Drosophila* imaginal discs by the Hedgehog receptor gene *patched*. Mech Dev 129:339–349.

27. Mast E, Bieser KL, Abraham-Villa M, Adams V, Akinlehin AJ, Aquino LZ, Austin JL, Austin AK, Beckham CN, Bengson EJ, Bieszk A, Bogard BL, Brennan RC, Brnot RM, Cirone NJ, Clark MR, Cooper BN, Cruz D, Daprizio KA, DeBoe J, Dencker MM, Donnelly LL, Driscoll L, DuBeau RJ, Durso SW, Ejub A, Elgosbi W, Estrada M, Evins K, Fox PD, France JM, Franco Hernandez MG, Garcia LA, Garl O, Gorsuch MR, Gorzeman-Mohr MA, Grothouse ME, Gubbels ME, Hakemiamjad R, Harvey CV, Hoeppner MA, Ivanov JL, Johnson VM, Johnson JL, Johnson A, Johnston K, Keller KR, Kennedy BT, Killian LR, Klumb M, Koehn OL, Koym AS, Kress KJ, Landis RE, Lewis KN, Lim E, Lopez IK, Lowe D, Luengo Carretero P, Lunaburg G, Mallinder SL, Marshall NA, Mathew J, Mathew J, Mcmanaway HS, Meegan EN, Meyst JD, Miller MJ, Minogue CK, Mohr AA, Moran CI, Moran A, Morris MD, Morrison MD, Moses EA, Mullins CJ, Neri CI, Nichols JM, Nickels BR, Okai AM, Okonmah C, Paramo M, Paramo M, Parker SL, Parmar NK, Paschal J, Patel P, Patel D, Perkins EB, Perry MM, Perry Z, Pollock AA, Portalatin O, Proffitt KS, Queen JT, Quemeneur AC, Richardson AG, Rosenberger K, Rutherford AM, Santos-Perez IX, Sarti CY, Schouweiler LJ, Sessing LM, Setaro SO, Silvestri CF, Smith OA, Smith MJ, Sumner JC, Sutton RR, Sweckard L, Talbott NB, Traxler PA, Truesdell J, Valenti AF, Verace L, Vijayakumar P, Wadley WL, Walter KE, Williams AR, Wilson TJ, Witbeck MA, Wobler TM, Wright LJ, Zuczkowska KA, Devergne O, Hamill DR, Shah HP, Siders J, Taylor EE, Vrailas-Mortimer AD, Kagey JD. 2022. Genetic mapping of *Uba3^O.2.2^*, a pupal lethal mutation in *Drosophila melanogaster*. MicroPublication Biol 2022.

28. Moore SL, Adamini FC, Coopes ES, Godoy D, Northington SJ, Stewart JM, Tillett RL, Bieser KL, Kagey JD. 2022. *Patched* and *Costal-2* mutations lead to differences in tissue overgrowth autonomy. Fly (Austin) 16:176–189.

29. Talley EM, Watts CT, Aboyer S, Adamson MG, Akoto HA, Altemus H, Avella PJ, Bailey R, Bell ER, Bell KL, Breneman K, Burkhart JS, Chanley LJ, Cook SS, DesLaurier MT, Dorsey TR, Doyle CJ, Egloff ME, Fasawe AS, Garcia KK, Graves NP, Gray TK, Gustafson EM, Hall MJ, Hayes JD, Holic LJ, Jarvis BA, Klos PS, Kritzmire S, Kuzovko L, Lainez E, McCoy S, Mierendorf JC, Neri NA, Neville CR, Osborn K, Parker K, Parks ME, Peck K, Pitt R, Platta ME, Powell B, Rodriguez K, Ruiz C, Schaefer MN, Shields AB, Smiley JB, Stauffer B, Straub D, Sweeney JL, Termine KM, Thomas B, Toth SD, Veile TR, Walker KS, Webster PN, Woodard BJ, Yoder QL, Young MK, Zeedyk ML, Ziegler LN, Bieser KL, Puthoff DP, Stamm J, Vrailas-Mortimer AD, Kagey JD, Merkle JA. 2021. Genetic mapping and phenotypic analysis of *shot^H.3.2^* in *Drosophila melanogaster*. MicroPublication Biol 2021.

30. Siders JL, Bieser KL, Hamill DR, Acosta EC, Alexander OK, Ali HI, Anderson MJ, Arrasmith HR, Azam M, Beeman NJ, Beydoun H, Bishop LJ, Blair MD, Bletch B, Bline HR, Brown JC, Burns KM, Calagua KC, Chafin L, Christy WA, Ciamacco C, Cizauskas H, Colwell CM, Courtright AR, Diaz Alavez L, Ecret RI, Edriss F, Ellerbrock TG, Ellis MM, Extine EM, Feldman E, Fickenworth LJ, Goeller CM, Grogg AS, Hernandez Y, Hershner A, Jauss MM, Jimenez Garcia L, Franks KE, Kazubski ET, Landis ER, Langub J, Lassek TN, Le TC, Lee JM, Levine DP, Lightfoot PJ, Love N, Maalhagh-Fard A, Maguire C, McGinnis BE, Mehta BV, Melendrez V, Mena ZE, Mendell S, Montiel-Garcia P, Murry AS, Newland RA, Nobles RM, Patel N, Patil Y, Pfister CL, Ramage V, Ray MR, Rodrigues J, Rodriquez VC, Romero Y, Scott AM, Shaba N, Sieg S, Silva K, Singh S, Spargo AJ, Spitnale SJ, Sweeden N, Tague L, Tavernini BM, Tran K, Tungol L, Vestal KA, Wetherbee A, Wright KM, Yeager AT, Zahid R, Kagey JD. 2021. Genetic Mapping of a new *Hippo* allele, *Hpo^N.1.2^*, in *Drosophila melanogaster*. MicroPublication Biol 2021.

31. Vrailas-Mortimer AD, Aggarwal N, Ahmed NN, Alberts IM, Alhawasli M, Aljerdi IA, Allen BM, Alnajar AM, Anderson MA, Armstong R, Avery CC, Avila EJ, Baker TN, Basardeh S, Bates NA, Beidas FN, Bosler AC, Brewer DM, Buenaventura RS, Burrell NJ, Cabrera-Lopez AP, Cervantes-Gonzalez AB, Cezar RP, Coronel J, Croslyn C, Damery KR, Diaz-Alavez L, Dixit NP, Duarte DL, Emke AR, English K, Eshun AA, Esterly SR, Estrada AJ, Feng M, Freund MM, Garcia N, Ghotra CS, Ghyasi H, Hale CS, Hulsman L, Jamerson L, Jones AK, Kuczynski M, Lacey-Kennedy TN, Lee MJ, Mahjoub T, Mersinger MC, Muckerheide AD, Myers DW, Nielsen K, Nosowicz PJ, Nunez JA, Ortiz AC, Patel TT, Perry NN, Poser WSA, Puga DM, Quam C, Quintana-Lopez P, Rennerfeldt P, Reyes NM, Rines IG, Roberts C, Robinson DB, Rossa KM, Ruhlmann GJ, Schmidt J, Sherwood JR, Shonoda DH, Soellner H, Torrez JC, Velide M, Weinzapfel Z, Ward AC, Bieser KL, Merkle JA, Stamm JC, Tillett RL, Kagey JD. 2021. *B.2.16* is a non-lethal modifier of the *Dark^82^* mosaic eye phenotype in *Drosophila melanogaster*. MicroPublication Biol 2021.

32. Bieser K, Sanford JS, Saville K, Arreola KF, Ayres ZT, Basulto D, Benito S, Breen CJ, Brix JA, Brown N, Burton KK, Chadwick TM, Chen M, Chu K, Corbett BL, Dill Z, Faughender MA, Hickey AD, Julia JS, Kelty SS, Kobs BBK, Krason BA, Lam B, McCullough CL, McEwen BR. 2019. Genetic mapping of *shn^E.3.2^* in *Drosophila melanogaster*. MicroPublication Biol https://doi.org/10.17912/micropub.biology.000118.

33. Bieser KL, Stamm J, Aldo AA, Bhaskara S, Claiborne M, Coronel Gómez JN, Dean R, Dowell A, Dowell E, Eissa M, Fawaz AA, Fouad-Meshriky MM, Godoy D, Gonzalez K, Hachem MK, Hammoud MF, Huffman A, Ingram H, Jackman AB, Karki B, Khalil N, Khalil H, Ha TK, Kharel A, Kobylarz I, Lomprey H, Lonnberg A, Mahbuba S, Massarani H, Minster M, Molina K, Molitor L, Murray T, Patel PM, Pechulis S, Raja A, Rastegari G, Reeves S, Sabu N, Salazar R, Schulert D, Senopole MD, Sportiello K, Torres C, Villalobos J, Wu J, Zeigler S, Kagey JD. 2018. The Mapping of *Drosophila melanogaster* mutant A.4.4. MicroPublication Biol.

34. Stamm J, Joshi G, Anderson MA, Bussing K, Houchin C, Elinsky A, Flyte J, Husseini N, Jarosz D, Johnson C, Johnson A, Jones C, Kooner T, Myhre D, Rafaill T, Sayed S, Swan K, Toma J, Kagey J. 2019. Genetic mapping of EgfrL.3.1 in Drosophila melanogaster. MicroPublication Biol 2019.

35. Evans CJ, Bieser KL, Acevedo-Vasquez KS, Augustine EJ, Bowen S, Casarez VA, Feliciano VI, Glazier A, Guinan HR, Hallman R, Haugan E, Hehr LA, Hunnicutt SN, Leifer I, Mauger M, Mauger M, Melendez NY, Milshteyn L, Moore E, Nguyen SA, Phanphouvong SC, Pinal DM, Pope HM, Salinas M-BM, Shellin M, Small I, Yeoh NC, Yokomizo AMK, Kagey JD. 2022. The *I.3.2* developmental mutant has a single nucleotide deletion in the gene *centromere identifier*. MicroPublication Biol 2022.

36. Cosenza A, Kagey JD. 2016. The Mapping and Characterization of *Cruella (Cru)*, a Novel Allele of *Capping Protein α (Cpa)*, Identified from a Conditional Screen for Negative Regulators of Cell Growth and Cell Division. Adv Biosci Biotechnol 07:373–380.

37. Ternovski J, Orr L, Kalla J, Aronow P. 2022. A Note on Increases in Inattentive Online Survey-Takers Since 2020. J Quant Descr Digit Media 2:1–35.

38. Little TD, Chang R, Gorrall BK, Waggenspack L, Fukuda E, Allen PJ, Noam GG. 2020. The retrospective pretest–posttest design redux: On its validity as an alternative to traditional pretest–posttest measurement. Int J Behav Dev 44:175–183.

39. Hake RR. 1998. Interactive-engagement versus traditional methods: A six-thousand-student survey of mechanics test data for introductory physics courses. Am J Phys 66:64–74.

40. Cook RK, Christensen SJ, Deal JA, Coburn RA, Deal ME, Gresens JM, Kaufman TC, Cook KR. 2012. The generation of chromosomal deletions to provide extensive coverage and subdivision of the *Drosophila melanogaster* genome. Genome Biol 13:R21, 1–14.

41. Hanauer DI, Graham MJ, Jacobs-Sera D, Garlena RA, Russell DA, Sivanathan V, Asai DJ, Hatfull GF. 2022. Broadening Access to STEM through the Community College: Investigating the Role of Course-Based Research Experiences (CREs). CBE—Life Sci Educ 21:ar38, 1–16.

42. Croonquist P, Falkenberg V, Minkovsky N, Sawa A, Skerritt M, Sustacek MK, Diotti R, Aragon AD, Mans T, Sherr GL, Ward C, Hall-Woods M, Goodman AL, Reed LK, Lopatto D. The Genomics Education Partnership: First findings on genomics research in community colleges. SPUR.

43. Bowman NA, Holmes JM. 2018. Getting off to a good start? First-year undergraduate research experiences and student outcomes. High Educ 76:17–33.

44. Neff LS, D’Souza MJ. 2019. Undergraduate Research, Data-Science Courses, and Volunteer Projects, Inform and Accelerate Wesley College’s Retention Among First- and Second-Year Students. Proc Natl Conf Undergrad Res Natl Conf Undergrad Res 2019:1434.

45. Hernandez PR, Woodcock A, Estrada M, Schultz PW. 2018. Undergraduate Research Experiences Broaden Diversity in the Scientific Workforce. BioScience 68:204–211.

46. Shaffer CD, Alvarez CJ, Bednarski AE, Dunbar D, Goodman AL, Reinke C, Rosenwald AG, Wolyniak MJ, Bailey C, Barnard D, Bazinet C, Beach DL, Bedard JEJ, Bhalla S, Braverman J, Burg M, Chandrasekaran V, Chung H-M, Clase K, DeJong RJ, DiAngelo JR, Du C, Eckdahl TT, Eisler H, Emerson JA, Frary A, Frohlich D, Gosser Y, Govind S, Haberman A, Hark AT, Hauser C, Hoogewerf A, Hoopes LLM, Howell CE, Johnson D, Jones CJ, Kadlec L, Kaehler M, Silver Key SC, Kleinschmit A, Kokan NP, Kopp O, Kuleck G, Leatherman J, Lopilato J, MacKinnon C, Martinez-Cruzado JC, McNeil G, Mel S, Mistry H, Nagengast A, Overvoorde P, Paetkau DW, Parrish S, Peterson CN, Preuss M, Reed LK, Revie D, Robic S, Roecklein-Canfield J, Rubin MR, Saville K, Schroeder S, Sharif K, Shaw M, Skuse G, Smith CD, Smith MA, Smith ST, Spana E, Spratt M, Sreenivasan A, Stamm J, Szauter P, Thompson JS, Wawersik M, Youngblom J, Zhou L, Mardis ER, Buhler J, Leung W, Lopatto D, Elgin SCR. 2014. A Course-Based Research Experience: How Benefits Change with Increased Investment in Instructional Time. CBE— Life Sci Educ 13:111–130.

47. Hewlett JA. 2018. Broadening Participation in Undergraduate Research Experiences (UREs): The Expanding Role of the Community College. CBE—Life Sci Educ 17:es9, 1–3.

