## Supplementary material for "Fly-CURE, a Multi-institutional CURE using *Drosophila*, Increases Students’ Confidence, Sense of Belonging, and Persistence in Research": Combined Appendices

### I. Fly-CURE institutional demographics

#### A. Demographic data of general student populations at Fly-CURE institutions

| Institution | State | Total student population | Under-graduate population | Undergrad -to-grad ratio | Female-to-male ratio | % minority | % Pell grant eligible | % First-generation |
|---|---|---|---|---|---|---|---|---|
| Albion College | MI | 1506 | 1506 | 1 | 1.15 | 35 | 47 | 25 |
| Anoka-Ramsey Community College | MN | 11213 | 11213 | 1 | 1.7 | 30 | 15 | 24 |
| Frostburg State University | MD | 4449 | 3677 | 0.826 | 1.27 | 46 | 34 | 26 |
| Illinois State University | IL | 20233 | 17674 | 0.874 | 1.5 | 27.4 | 32 | N/A |
| Loyola Marymount University | CA | 9686 | 6096 | 0.629 | 1.17 | 47.1 | N/A | 10 |
| Morehouse College | GA | 2554 | 2554 | 1 | 0 | 99 | 57 | N/A |
| Nevada State College | NV | 7215 | 7201 | 0.998 | 2.8 | 67 | 38.4 | 29.4 |
| Northern Illinois University | IL | 16234 | 11834 | 0.729 | 1.26 | 52.65 | 50.9 | 49.6 |
| Ohio Northern University | OH | 3116 | 2191 | 0.703 | 0.92 | 11 | 21 | 16 |
| Ohio Wesleyan University | OH | 1426 | 1426 | 1 | 1.24 | 22.6 | 28 | 18.1 |
| The College of Wooster | OH | 1973 | 1973 | 1 | 1.19 | 22.4 | N/A | N/A |
| University of Detroit Mercy | MI | 5227 | 2924 | 0.559 | 1.61 | 19 | 28 | 33 |
| University of Evansville | IN | 2078 | 1796 | 0.864 | 1.5 | 15 | 25 | 16 |
| University of St. Francis | IL | 3426 | 1316 | 0.384 | 2.03 | 46 | 43 | 46 |
| Western New Mexico University | NM | 3632 | 2848 | 0.784 | 2 | 51 | 62 | 54 |

N/A = Not available

#### B. Classifications and designations of Fly-CURE institutions

| Institution | Specific Carnegie classification | General Carnegie classification | Minority Serving Institution (MSI) category |
|---|---|---|---|
| Albion College | Baccalaureate Colleges: Arts & Sciences Focus | PUI | no |
| Anoka-Ramsey Community College | Associate's Colleges: High Transfer-High Nontraditional | Community college | no |
| Frostburg State University | Master's Colleges & Universities: Larger Programs | Masters university | no |
| Illinois State University | Doctoral Universities: High Research Activity | R2 | no |
| Loyola Marymount University | Doctoral Universities: High Research Activity | R2 | no |
| Morehouse College | Baccalaureate Colleges: Arts & Sciences Focus | PUI | HBCU |
| Nevada State College | Baccalaureate Colleges: Diverse Fields | PUI | AANAPISI and HSI |
| Northern Illinois University | Doctoral Universities: High Research Activity | R2 | no |
| Ohio Northern University | Baccalaureate Colleges: Diverse Fields | PUI | no |
| Ohio Wesleyan University | Baccalaureate Colleges: Arts & Sciences Focus | PUI | no |
| The College of Wooster | Baccalaureate Colleges: Arts & Sciences Focus | PUI | no |
| University of Detroit Mercy | Doctoral/Professional Universities | Doctoral/Professional university | no |
| University of Evansville | Master's Colleges & Universities: Medium Programs | Masters university | no |
| University of St. Francis | Doctoral/Professional Universities | Doctoral/Professional university | no |
| Western New Mexico University | Master's Colleges & Universities: Larger Programs | Masters university | HSI |

AANAPISI = Asian American and Native American Pacific Islander-Serving Institution; HBCU = Historically Black College or University; HSI = Hispanic Serving Institution; MSI = Minority Serving Institution; PUI = Primarily Undergraduate Institution

#### C. Fly-CURE modules and associated courses taught at participating institutions

| Institution | Course | Components included in course | | | |
|---|---|---|---|---|---|
| | | Complementation mapping | Phenotypic characterization | Molecular analysis | Bioinformatics |
| Albion College | Genetics | Y | Y | Y | Y |
| Anoka-Ramsey Community College | Genetics | Y | Y | Y | Y |
| Frostburg State University | Genetics | Y | Y | Y | Y |
| Illinois State University | Genetics | Y | Y | Y | Y |
| Loyola Marymount University | Genetics Laboratory | Y | Y | Y | Y |
| Morehouse College | Genetics | Y | Y | N/A | Y |
| Nevada State College | Genetics | Y | Y | Y | Y |
| Northern Illinois University | Genetics | Y | Y | Y | N |
| Ohio Northern University | Genetics | Y | Y | Y | Y |
| Ohio Wesleyan University | Genetics | Y | Y | Y | Y |
| The College of Wooster | Molecular Biology | Y | Y | Y | Y |
| University of Detroit Mercy | Genetics Laboratory | Y | Y | Y | Y |
| University of Evansville | Genetics | Y | Y | Y | Y |
| University of St. Francis | Introductory Biology | Y | Y | Y | Y |
| Western New Mexico University | Anatomy and Physiology | N | Y | N | N |

Y = Yes; N = No; N/A = not available

#### II. Fly-CURE student demographics

##### A. Gender

| | | Frequency | Percent | Valid Percent |
|---|---|---|---|---|
| Valid | Male | 130 | 27 | 28 |
| | Female | 318 | 66 | 69 |
| | Not listed | 4 | 1 | 1 |
| | Prefer not to say | 8 | 2 | 2 |
| | Total | 460 | 96 | 100 |
| Missing | Blank | 20 | 4 | |
| Total | | 480 | 100 | |

##### B. Academic year

| | | Frequency | Percent | Valid Percent |
|---|---|---|---|---|
| Valid | Freshman | 20 | 4 | 4 |
| | Sophomore | 156 | 33 | 34 |
| | Junior | 143 | 30 | 31 |
| | Senior | 132 | 28 | 29 |
| | Already have a bachelor's degree | 9 | 2 | 2 |
| | Total | 460 | 96 | 100 |
| Missing | Blank | 20 | 4 | |
| Total | | 480 | 100 | |

##### C. Student rank (combined)

| | | Frequency | Percent | Valid Percent |
|---|---|---|---|---|
| Valid | Freshman/Sophomore | 176 | 37 | 38 |
| | Junior/Senior/Already has bachelor's degree | 284 | 59 | 62 |
| | Total | 460 | 96 | 100 |
| Missing | Blank | 20 | 4 | |
| Total | | 480 | 100 | |

##### D. Underrepresented in STEM

| | | Frequency | Percent | Valid Percent |
|---|---|---|---|---|
| Valid | Not Underrepresented in STEM | 335 | 70 | 73 |
| | Underrepresented in STEM | 124 | 26 | 27 |
| | Total | 459 | 96 | 100 |
| Missing | Blank | 21 | 4 | |
| Total | | 480 | 100 | |

##### E. First generation student status

| | | Frequency | Percent | Valid Percent |
|---|---|---|---|---|
| Valid | First generation student (neither mother nor father attended college) | 134 | 28 | 29 |
| | Continued generation student (one or both parents attended college) | 323 | 67 | 71 |
| | Total | 457 | 95 | 100 |
| Missing | Blank | 23 | 5 | |
| Total | | 480 | 100 | |

#### F. Previous research experience

| | | Frequency | Percent | Valid Percent |
|---|---|---|---|---|
| Valid | No research exposure | 228 | 48 | 48 |
| | Some research exposure | 252 | 53 | 53 |
| | Total | 480 | 100 | 100 |

#### G. Types of prior research experiences

| | Yes | | No | | Total |
|---|---|---|---|---|---|
| | Frequency | Percent | Frequency | Percent | |
| a. Took one or more Introduction to Research OR Research Methods courses | 147 | 31 | 332 | 69 | 479 |
| b. Took a course where the outcome of the research was not already known | 142 | 30 | 336 | 70 | 478 |
| c. Did research during the summer with a research mentor (PI or grad student) | 66 | 14 | 413 | 86 | 479 |
| d. Did research during the semester with a research mentor | 126 | 26 | 354 | 74 | 480 |

**Fly-CURE Course Pre-Survey** [selected questions]

1. Are you: (check all that apply)
   - Native Hawaiian or other Pacific Islander (original peoples)
   - American Indian or Alaskan Native
   - Asian (including subcontinent and Philippines)
   - Black or African American (including African and Caribbean)
   - White

2. Are you Hispanic or Latino?
   - Yes
   - No

3. Are you of Middle Eastern descent (including Arab states, Israel, Turkey, Afghanistan, Iran, Pakistan, Egypt, Iraq, and Saudi Arabia)?
   - Yes
   - No

4. What is your gender?
   - Male
   - Female
   - Not listed
   - Prefer not to say

5. What is your current academic standing?
   - Freshman
   - Sophomore
   - Junior
   - Senior
   - Already have my bachelor's degree

6. Has either your mother or father attended any college?
   - Both mother and father attended college
   - Only mother attended college
   - Only father attended college
   - Neither mother nor father attended college

7. Which of these previous research experiences have you had?
   a. Took one or more Introduction to Research <u>or</u> Research Methods courses?
      i. Yes
      ii. No
   b. Took a course where the outcome of the research was not already known?
      i. Yes
      ii. No
   c. Did research during the <u>summer</u> with a research mentor (PI or grad student)?
      i. Yes
      ii. No
   d. Did research during the <u>semester</u> with a research mentor?
      i. Yes
      ii. No
   e. Presented my research to <u>others at my university</u>?

i. Yes
ii. No

f. Presented my research <u>outside</u> of my university?
   i. Yes
   ii. No
g. Had some other kind of research experience prior to this course?
   i. Yes (please describe this experience below)
   ii. No

8. Please tell us how confident you feel, now, about each of these…

| | 1 – Not at all | 2 | 3 | 4 | 5 – Completely |
|---|---|---|---|---|---|
| I can use research tools, instruments, or techniques to measure concepts of interest | | | | | |
| I can generate a research question to answer | | | | | |
| I can figure out what data/observations to collect and how to collect them | | | | | |
| I can explain the results of a study | | | | | |
| I can use scientific literature and/or reports to guide research | | | | | |
| I can integrate results from multiple studies into a theme or theory | | | | | |
| I can solve problems through research | | | | | |
| I can explain my scientific research to others | | | | | |

9. Please tell us how strongly you agree or disagree with each of these…

| | Strongly disagree | Disagree | Neutral | Agree | Strongly agree |
|---|---|---|---|---|---|
| I feel I belong to a community of researchers | | | | | |
| I have come to think of myself as a ‘researcher’ | | | | | |
| I feel like I belong in the field of research | | | | | |
| I gain satisfaction from doing important research | | | | | |
| Please mark "neutral" for this item | | | | | |

**Fly-CURE Course Post-Survey** [selected questions]

1. How interested are you in taking another science lab course that incorporates research where the outcome is not known?
   - Very interested
   - Somewhat interested
   - Not interested at all

2. How likely are you to:

| | Not likely | A little likely | Somewhat likely | Very likely | Definitely |
|---|---|---|---|---|---|
| a) Enroll in another research intensive science lab course? | | | | | |
| b) Pursue or continue independent research in a science lab? | | | | | |
| c) Pursue a career as a scientist? | | | | | |

3. Compared to your intentions **BEFORE** taking this course, **HOW LIKELY ARE YOU NOW** to:

| | Not more likely | A little more likely | Somewhat more likely | Much more likely | Extremely more likely |
|---|---|---|---|---|---|
| a) Enroll in another research intensive science lab course? | | | | | |
| b) Pursue or continue independent research in a science lab? | | | | | |
| c) Pursue a career as a scientist? | | | | | |

4. Please tell us how confident you feel, now, about each of these…

| | 1 – Not at all | 2 | 3 | 4 | 5 – Completely |
|---|---|---|---|---|---|
| I can use research tools, instruments, or techniques to measure concepts of interest | | | | | |
| I can generate a research question to answer | | | | | |
| I can figure out what data/observations to collect and how to collect them | | | | | |
| I can explain the results of a study | | | | | |
| I can use scientific literature and/or reports to guide research | | | | | |

| |
|---|
| I can integrate results from multiple studies into a theme or theory |
| I can solve problems through research |
| I can explain my scientific research to others |

5. Please tell us how strongly you agree or disagree with each of these…

| | Strongly disagree | Disagree | Neutral | Agree | Strongly agree |
|---|---|---|---|---|---|
| I feel I belong to a community of researchers | | | | | |
| I have come to think of myself as a 'researcher' | | | | | |
| I feel like I belong in the field of research | | | | | |
| I gain satisfaction from doing important research | | | | | |
| Please mark "neutral" for this item | | | | | |
