## Supplemental Figures for "Fly-CURE, a Multi-institutional CURE using *Drosophila*, Increases Students’ Confidence, Sense of Belonging, and Persistence in Research"

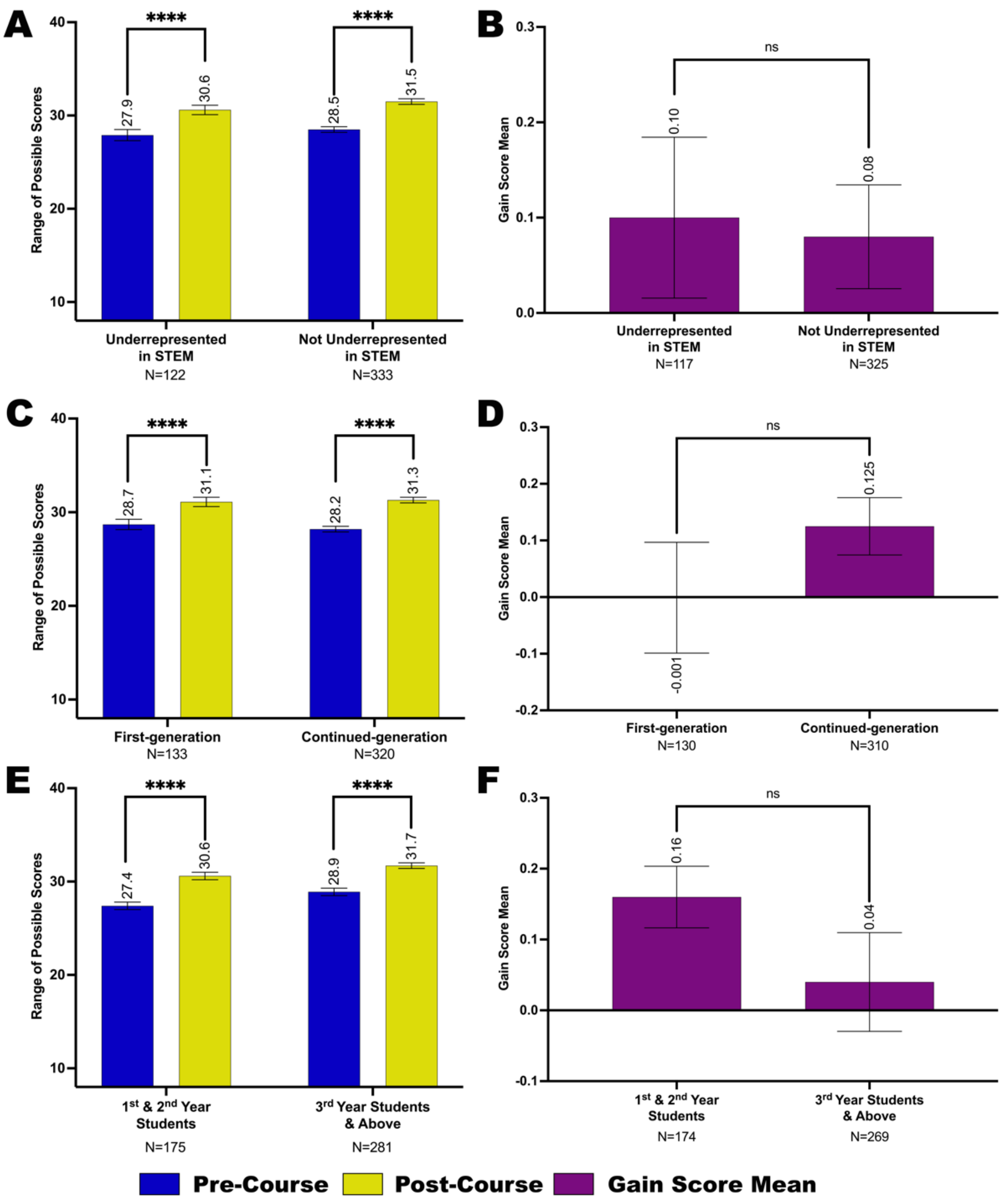


**Figure S1. Reported self-efficacy in research for student subgroups before and after completing the Fly-CURE.** The mean self-efficacy pre-course (blue) and post-course (yellow) scale score means are shown for participant subgroups, as well as the gain score mean (purple) to compare differential gains in self-efficacy post-course compared to pre-course in student subgroups. (A-B) Self-efficacy scale score means (A) and gain score mean (B) for minority students underrepresented in STEM and students not considered underrepresented in STEM. (C-D) Self-efficacy scale score mean (C) and gain score mean (D) for first-generation and continued-generation college students. (E-F) Self-efficacy scale score mean (E) and gain score mean (F) for first- or second-year students compared to third-year students and above. Error bars, ±SEM; ns, not significant, $P > 0.05$; ****$P \leq 0.0001$.

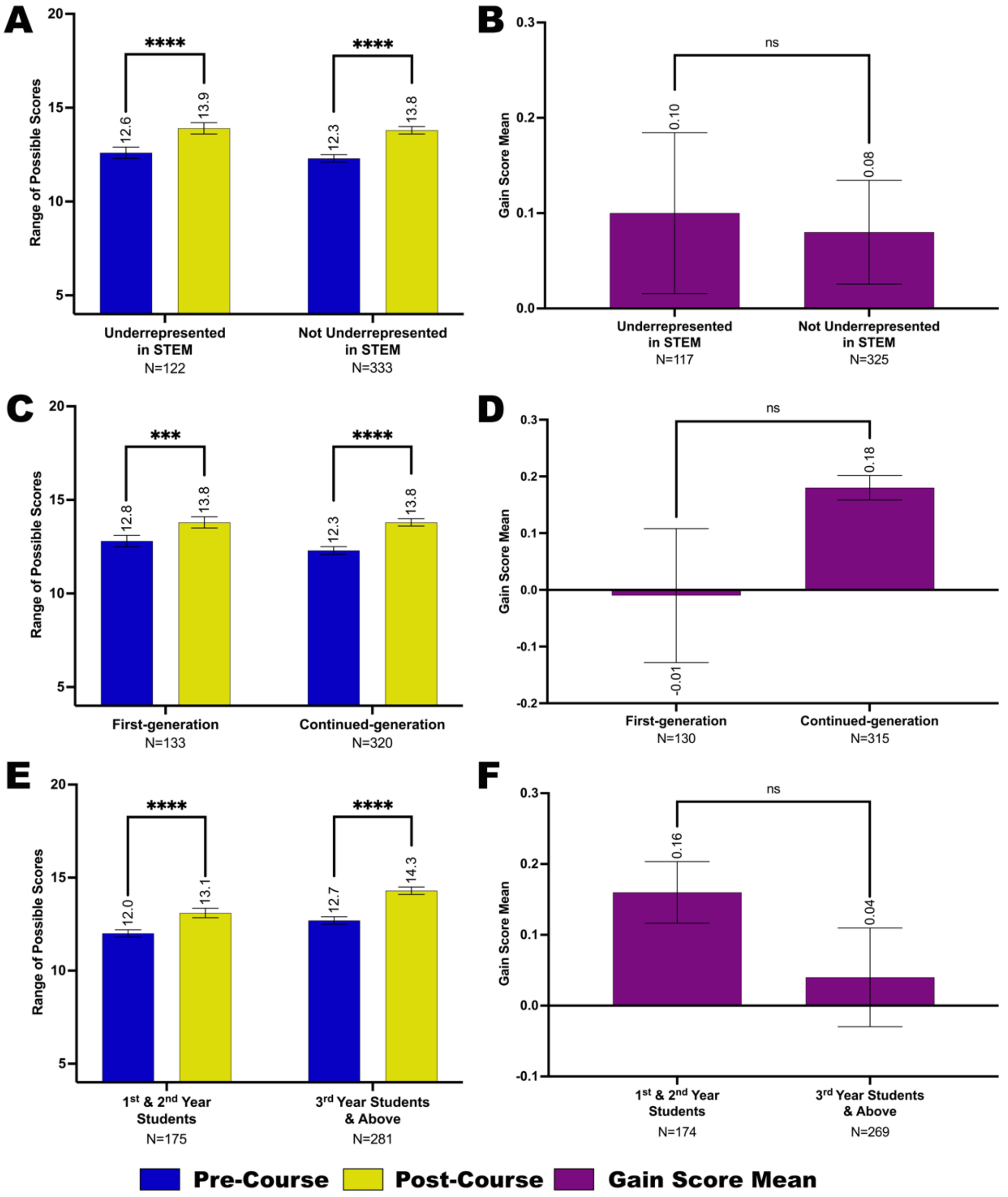


**Figure S2. Reported sense of belonging in science for student subgroups before and after completing the Fly-CURE.** The scale score means for sense of belonging in the scientific community pre-course (blue) and post-course (yellow) are shown for participant subgroups, as well as the gain score mean (purple) to compare differential gains in sense of belonging post-course compared to pre-course in student subgroups. (A-B) Sense of belonging scale score mean (A) and gain score mean (B) for minority students underrepresented in STEM and students not considered underrepresented in STEM. (C-D) Sense of belonging in research scale score mean (C) and gain score mean (D) for first-generation and continued-generation college students. (E-F) Sense of belonging scale score mean (E) and gain score mean (F) for first- or second-year students compared to third-year students and above. Error bars, ±SEM; ns, not significant, $P > 0.05$; $***P \leq 0.001$; $****P \leq 0.0001$.

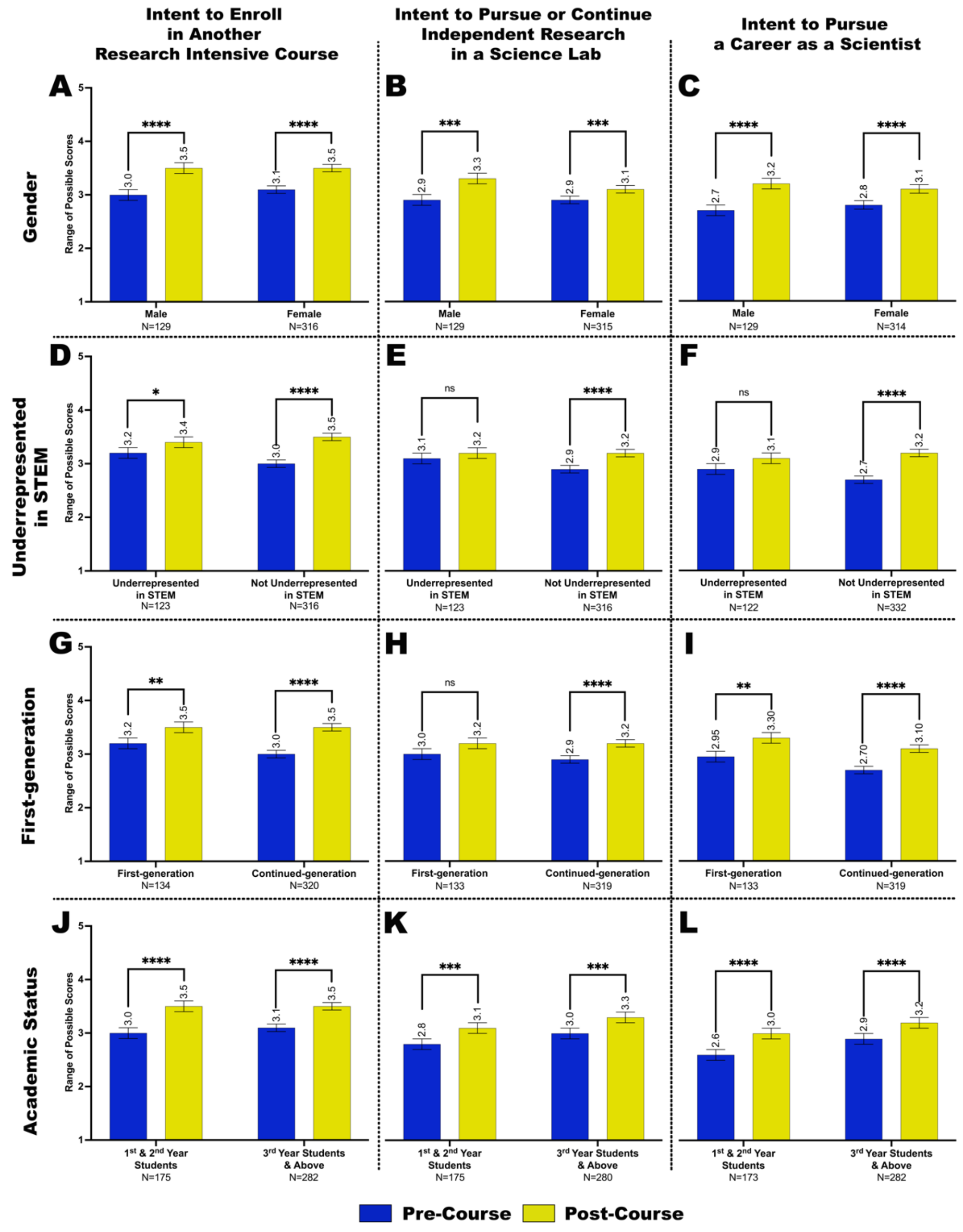


**Figure S3. Reported intent to seek additional research experiences for student subgroups before and after completing the Fly-CURE.** Comparison of students' perceived interest before (blue) and after (yellow) the CURE to enroll in another research-intensive science laboratory course (A,D,G,J), pursue or continue independent research in a research laboratory (B,E,H,K), and pursue a career as a scientist (C,F,I,L). Scale score means for reported perceived interest in seeking additional research experiences before compared to after the Fly-CURE for male and female students (A-C), minority students underrepresented in STEM and students not considered underrepresented in STEM (D-F), first-generation and continued-generation college students (G-I), and first- or second-year students compared to third-year students and above (J-L). Error bars, ±SEM; ns, not significant, $P > 0.05$; $*P \leq 0.05$; $**P \leq 0.01$; $***P \leq 0.001$; $****P \leq 0.0001$.

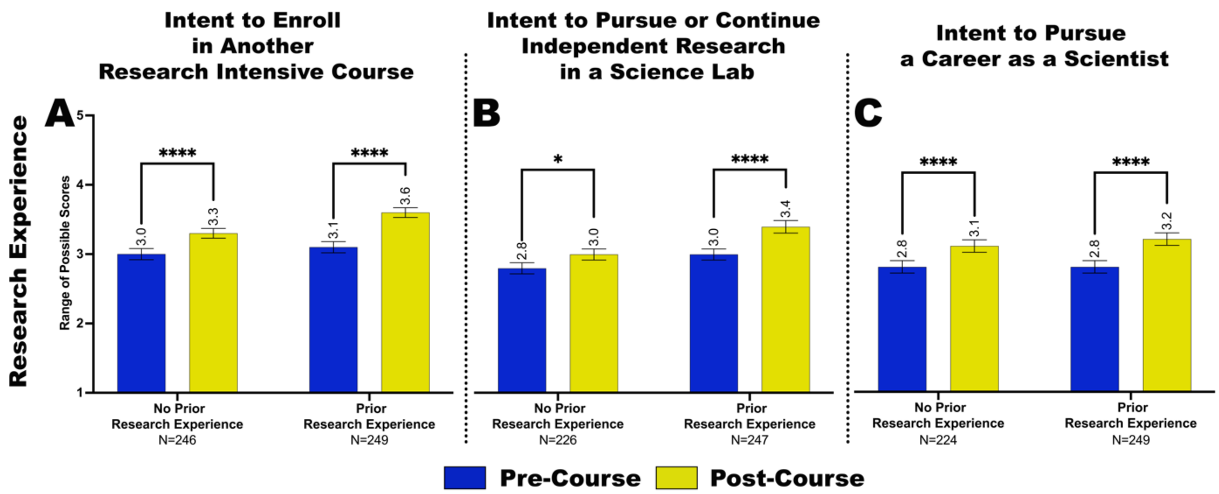


**Figure S4. Reported intent to seek additional research experiences in students with and without research experience prior to the Fly-CURE.** Through post-surveys, students reported their perceived interest in pursuing additional research–associated experiences before and after completing the Fly-CURE. The survey rating scales ranged from one (not likely) to five (definitely) for the research experiences indicated. Scale score means of perceived student interest to enroll in another research-intensive science laboratory course (A), pursue or continue independent research in a research laboratory (B), and pursue a career as a scientist (C) before (blue) and after (yellow) taking the Fly-CURE for students who reported as having or not having research experience prior to the Fly-CURE. Error bars, ±SEM; *$P \leq 0.05$; ****$P \leq 0.0001$.
